# Humanized antibody potently neutralizes all SARS-CoV-2 variants by a novel mechanism

**DOI:** 10.1101/2022.06.26.497634

**Authors:** Sai Luo, Jun Zhang, Alex J.B. Kreutzberger, Amanda Eaton, Robert J. Edwards, Changbin Jing, Hai-Qiang Dai, Gregory D. Sempowski, Kenneth Cronin, Robert Parks, Adam Yongxin Ye, Katayoun Mansouri, Maggie Barr, Novalia Pishesha, Aimee Chapdelaine Williams, Lucas Vieira Francisco, Anand Saminathan, Hanqin Peng, Himanshu Batra, Lorenza Bellusci, Surender Khurana, S. Munir Alam, David C. Montefiori, Kevin O. Saunders, Ming Tian, Hidde Ploegh, Tom Kirchhausen, Bing Chen, Barton F. Haynes, Frederick W. Alt

## Abstract

SARS-CoV-2 Omicron variants have generated a world-wide health crisis due to resistance to most approved SARS-CoV-2 neutralizing antibodies and evasion of antibodies induced by vaccination. Here, we describe the SARS-CoV-2 neutralizing SP1-77 antibody that was generated from a humanized mouse model with a single human V_H_1-2 and Vκ1-33-associated with immensely diverse complementarity-determining-region-3 (CDR3) sequences. SP1-77 potently and broadly neutralizes SARS-CoV-2 variants of concern and binds the SARS-CoV-2 spike protein receptor-binding-domain (RBD) via a novel-CDR3-based mode. SP1-77 does not block RBD-binding to the ACE2-receptor or endocytosis step of viral entry, but rather blocks membrane fusion. Our findings provide the first mechanistic insight into how a non-ACE2 blocking antibody potently neutralizes SARS-CoV-2, which may inform strategies for designing vaccines that robustly neutralize current and future SARS-CoV-2 variants.

## MAIN TEXT

The SARS-CoV-2 Omicron variant of concern has spread worldwide and remains the cause of a public health crisis. This variant has an unprecedented number of mutations in its spike protein, is resistant to most prior SARS-CoV-2-neutralizing antibodies, and evades antibodies induced by vaccinations (*1–6*). The Omicron BA.2 sub-variant dominates cases in South Africa, China, and other countries and has exacerbated the impact of the Omicron wave world-wide. Moreover, new Omicron sub-variants continue to emerge, including BA2.12.1, BA.4 and BA.5 (*7–9*). The few published human monoclonal antibodies that neutralize omicron BA.1 and BA.2 came from B cells of human patients previously infected by SARS-CoV-2 (*1–6*). To manage Omicron sub-variants and prepare for potential future SARS-CoV-2 variants of concern (VOCs), new human or humanized antibodies that robustly neutralize all SARS-CoV-2 VOCs are needed. In addition, a characterization of their mechanism of action will be essential.

Binding of the SARS-CoV-2 spike protein to its obligate angiotensin-converting enzyme 2 (ACE2) receptor on the target cell surface initiates infection (*10*). The spike protein is made up of non-covalently linked S1 and S2 subunits (*11*). The receptor-binding-domain (RBD) for ACE2 is located in the S1 subunit, while the S2 subunit anchors the spike protein in the viral membrane. The S2 subunit also contains other sequences that mediate viral/host cell membrane fusion for viral entry. The RBD can adopt two distinct conformations: “down” and “up”; the “down” state is shielded from ACE2 binding, while the “up” state is ACE2-accessible (*12*). Engagement of ACE2 by an “up” RBD exposes the S2’ cleavage site on the S2 subunit to either the serine 2 (TMPRSS2) transmembrane protease at the infected cell surface, or to cathepsin L in the endosomal compartment following endocytosis of the ACE2/SARS-CoV-2 complex (*13*). Cleavage of the S2 subunit by these proteases leads to dissociation of the S1 subunit, which exposes the fusion peptide on the S2 subunit and ultimately leads to fusion pore formation, viral-host membrane fusion, and viral entry into the infected cell (*14*).

Therapeutic SARS-CoV-2 neutralizing antibodies (nAbs) that block virus entry into host cells have demonstrated substantial efficacy for treating COVID-19 infections. The majority of such nAbs target the SARS-CoV-2 RBD and inhibit viral entry by binding to the ACE2 receptor binding motif (RBM), thereby, directly impeding its binding to the ACE2 receptor(*1*). Other nAbs bind outside of the RBM, but sterically inhibit ACE2 binding (*15*). A few reported SARS-CoV-2 nAbs bind outside of the RBM and do not inhibit ACE2 binding, but can still potently neutralize SARS-CoV-2 VOCs (*1, 16, 17*). How the latter class of nAbs neutralizes SARS-CoV-2 is not known.

A vast portion of the primary antibody repertoire diversity lies in the antigen-contact CDR3-encoding variable region sequences generated *de novo* by non-templated V(D)J junctional modifications during V(D)J recombination (*18*). Indeed, the number of CDR3 sequences that can be generated through junctional diversification exceeds the number of B cells in mammalian immune systems by many orders of magnitude (*19*). Thus, the relative size of the primary BCR repertoire in humans and mice is determined by the number of primary B cells in their immune system (*20*). Immunoglobin (Ig) gene-humanized mice capable of rearranging the full complement of human *Igh* and *IgL* variable region gene segments have yielded many therapeutic monoclonal antibodies (mAbs), including mAbs that neutralize several SARS-CoV-2 VOCs (*15*). However, these mouse models have not yet been reported to yield Omicron-neutralizing mAbs. There could be many possible explanations for the lack of isolation of mAbs that more broadly neutralize SARS-CoV-2 VOCs from Immunoglobin (Ig) gene-humanized mice. One of these possibilities is that mice express only a tiny fraction of the CDR3-diversity present in human primary BCR repertoire, as a mouse has 1000-fold fewer B cells than a human. Early experiments indicated that BCR repertoires generated from rearrangement of a single V_H_ and Vκ could make potent responses to diverse antigens due to immense CDR3 diversification (*21*). Building from this observation, we now report that a mouse model that generates more human-like CDR3 diversity in association with rearrangement of a single human V_H_ and Vκ can indeed be used to elicit novel SARS-CoV-2 nAbs.

## RESULTS

### Single human V_H_- and Vκ-rearranging mouse model for human antibody discovery

For these studies, we generated the “V_H_1-2/Vκ1-33-rearranging mouse model” that generates primary BCR repertoires via exclusive rearrangement of a single human V_H_1-2 and predominant rearrangement of human Vκ1-33 with the primary humanized BCR repertoire diversification based on immense CDR3 diversification (Fig.1A). In this regard, V_H_1-2 is frequently represented in SARS-CoV-2 nAbs derived from human patients, but has not contributed to antibodies that broadly neutralize the different VOCs (*22*). For the *Igh* locus in this model, the proximal mouse V_H_5-2 was replaced with human V_H_1-2, which linked human V_H_1-2 to the mouse V_H_5-2 downstream CTCF-binding element (CBE) (*23, 24*). All functional mouse V_H_s were deleted from this allele, and robust V_H_1-2 rearrangement, mediated by the linked CBE, was enforced through deletion of IGCR1 element in the V_H_ to D interval (*23–25*) (Fig.1A and fig. S1A). For the *Igκ* locus in this model, the proximal mouse Vκ3-2 that expressed in 2.7% of splenic B cells was replaced with human Vκ1-33, which resulted in expression in 2.2% of splenic B cells (Fig.1, A and B; fig. S1, B and C; table S1). Then, the IGCR1-related Cer/Sis element in the Vκ-Jκ interval was deleted to substantially increase Vκ1-33 utilization to 11% (*26*) (Fig.1, A and B; fig. S1, D and E; table S1). We retained the mouse J_H_s and Jκs in this model as their framework sequences outside of CDR3 are largely conserved between mouse and human (fig. S1F). We also retained the mouse Ds, as mouse Ds are highly related to a subset of human Ds and, in any case, contribution of D segments to CDR3 are often unassignable due to extensive non-templated junctional diversification.

**Figure 1.**
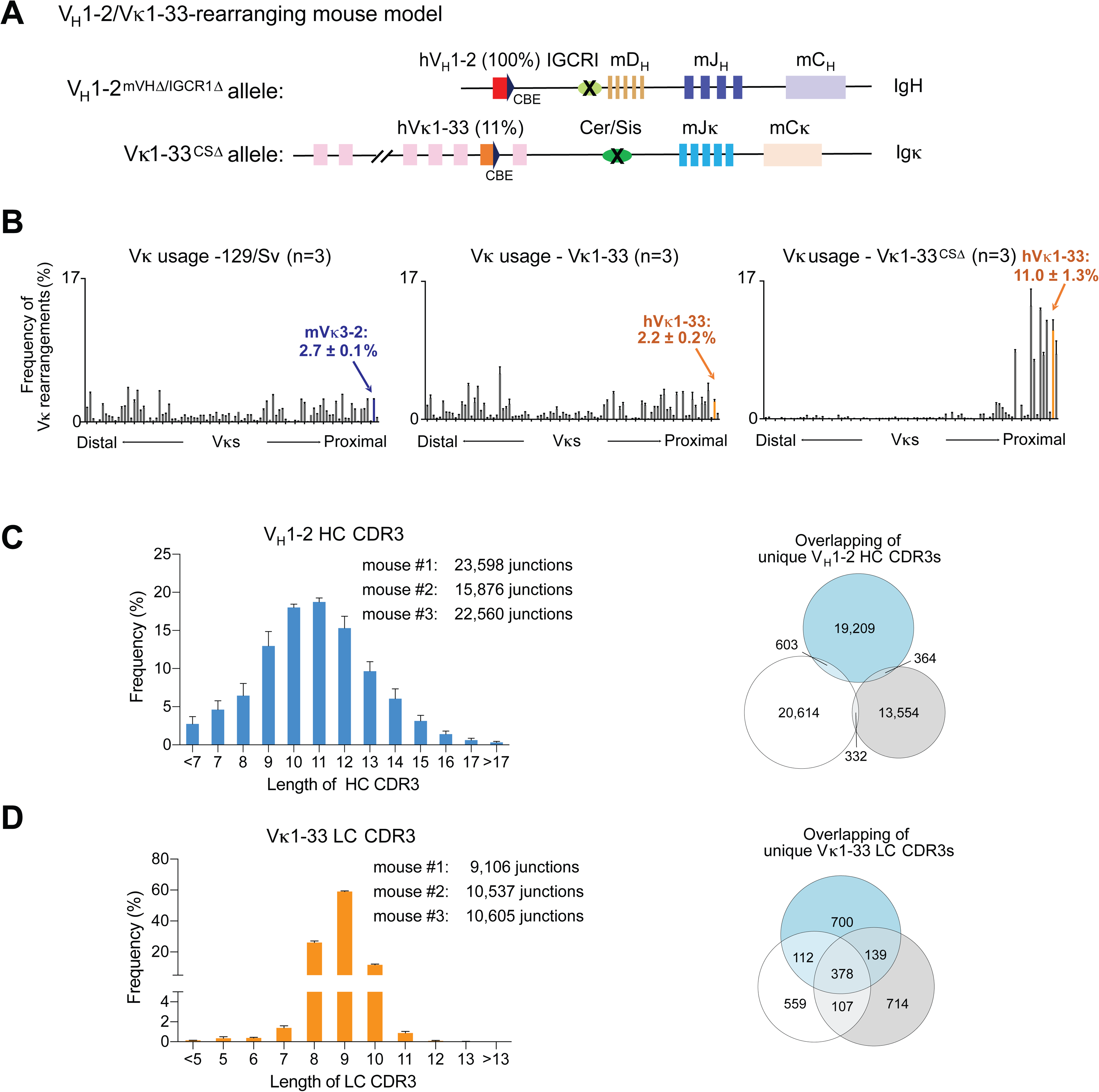
A humanized mouse model with diverse BCR repertoire derived from single human V_H_ and Vκ recombination. (**A**) Schematic representation of genetic modifications in the *Igh* and *Igκ* locus of V_H_1-2/Vκ1-33-rearranging mouse model. The V_H_1-2^IGCR1Δ^ allele was made previously (*23*). We deleted 2MB region upstream of V_H_1-2 that contains all the mouse V_H_s to generate V_H_1-2^mVHΔ/IGCR1Δ^ allele. We replaced mouse Vκ3-2 to human Vκ1-33 and deleted the Cer/sis element in Vκ to Jκ interval to generate Vκ1-33^CSΔ^ allele. (**B**) HTGTS-rep-seq analysis of Vκ usages in 129/Sv wild-type (left panel) and Vκ1-33-rearranging mouse splenic B cells in presence (middle panel) and absence (right panel) of Cer/sis element. The x axis lists all functional Vκs from the distal to the *Jκ*-proximal end. The histogram displays the percent usage of each Vκ among all productive VκJκ rearrangements. The junction number of each Vκ was shown in table S1. (**C-D**) The length distribution of V_H_1-2 HC CDR3 (left panel in (**C**)) and Vκ1-33 LC CDR3 (left panel in (**D**)) in splenic B cells. Data are mean ± SD of three libraries from different mice. Venn diagram shows the V_H_1-2 HC CDR3 (right panel in (**C**)) and Vκ1-33 LC CDR3 (left panel in (**D**)) complexity. The unique reads derived from the same libraries in the left panel. Little overlap of V_H_1-2 HC CDR3 sequences among three independent mice indicates tremendous CDR3 complexity.

In the resulting V_H_1-2/Vκ1-33-rearranging mouse model, human V_H_1-2 and Vκ1-33 contribute to the BCRs expressed by 100% and 11%, respectively, of splenic B cells (Fig. 1B, fig. S2A and table S1). Splenic B and T cell populations in this mouse model were similar in numbers and characteristics to those of wild-type as assessed by cytofluorimetry (fig. S2B). Moreover, analysis of CDR3s in primary BCR repertoires of these mice revealed an immensely diverse V_H_1-2-based CDR3 repertoire, as well as a diverse Vκ1-33 repertoire (Fig. 1, C and D). Following immunization, we predicted that the massive CDR3 diversity would contribute to selection into germinal centers of B cells with human V_H_1-2- and Vκ1-33-based BCRs, where somatic hypermutations could further diversify variable region exons and contribute to affinity maturation.

### Identification of novel SARS-CoV-2 neutralizing antibodies

We immunized the human V_H_1-2/Vκ1-33-rearranging model with recombinant stabilized soluble SARS-CoV-2 Wuhan-1 spike trimer, or with an RBD monomer fused to a nanobody (“VHH7-RBD”) that targets MHC II-complex antigens (*27*) to potentially increase immunogenicity--given the poor immunogenicity of RBD monomer (*28*) (Fig. 2A). Immunizations were done twice, four weeks apart, with each immunogen plus poly I:C adjuvant (Fig. 2A). All mice developed strong antibody responses to the SARS-CoV-2 spike or RBD immunogens, as demonstrated by serum IgG titers at 2 weeks and 6 weeks post immunization (fig. S3A). At 3 weeks after the second immunization, 96 antigen-specific IgG^+^ B cells were fluorescence-activated single-cell sorted from each mouse and their IgH (HC) and IgL (LC) variable region exons were sequenced (fig. S3B). These experiments identified 9 separate spike-specific B cell lineages, each of which expressed BCRs with unique sets of V_H_1-2 and Vκ1-33-associated CDR3s (Fig. 2B). In each CDR3-based lineage, individual members had unique patterns of somatic hypermutations (table. S2). The V_H_1-2-based HC CDR3s of these lineages were also unique compared to CDR3s of previously reported V_H_1-2-based anti-SARS-CoV-2 antibodies. To assess binding properties, we expressed one mAb from each lineage with human IgG1or IgG4 and human Igκ constant regions. ELISAs showed that all bind to the SARS-CoV-2 spike protein and 6 bind to the RBD, with one of the latter, termed SP1-77, derived from spike protein immunization (fig. S3C).

**Figure 2.**
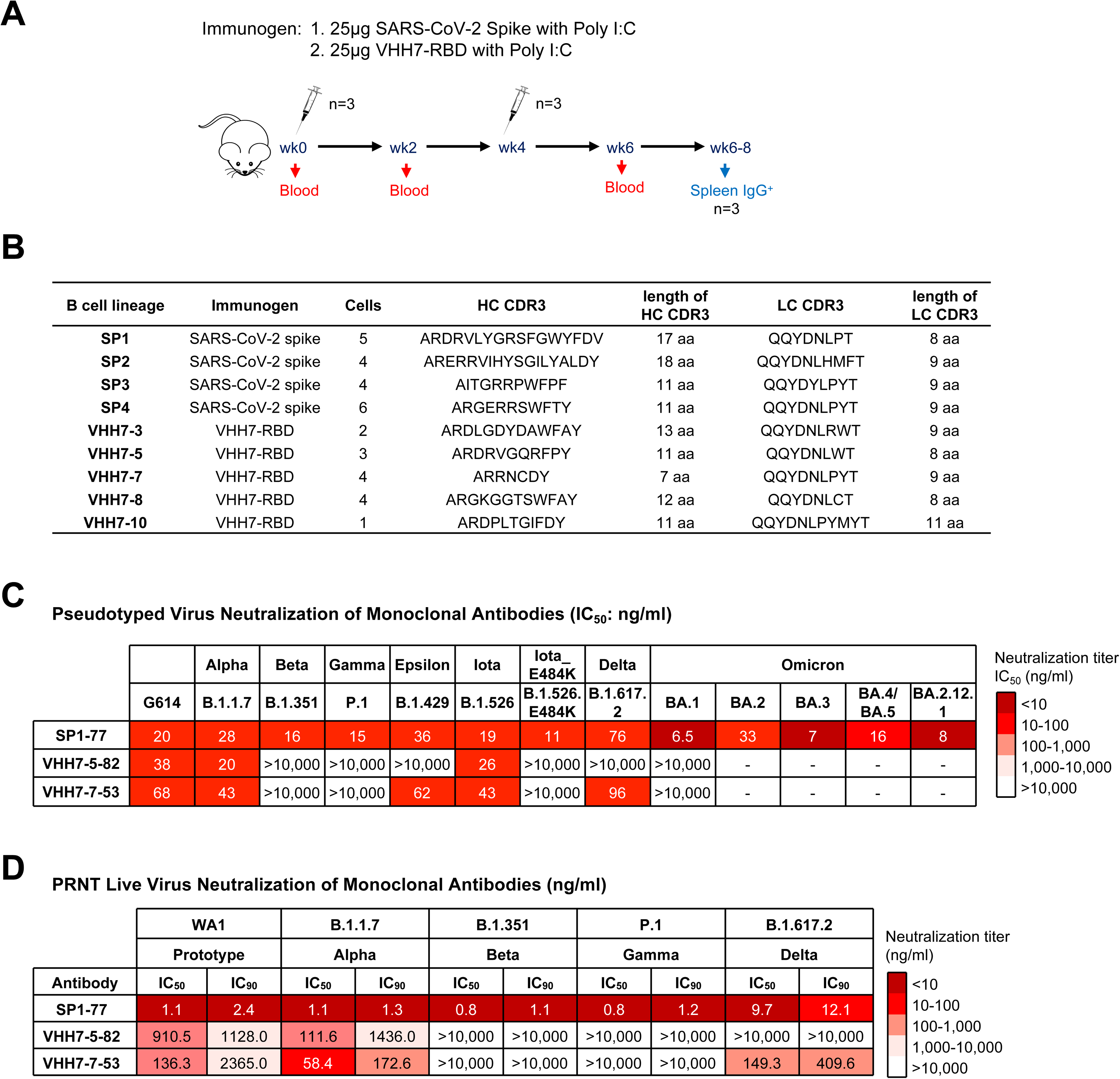
Immunizing the mouse model with SARS-CoV-2 spike or RBD elicited potent and broad V_H_1-2/Vκ1-33 antibodies. (**A**) Immunization scheme. Prime plus boost immunizations were performed at an interval of four weeks. (**B**) Table shows the V_H_1-2/Vκ1-33 antibodies isolated from SARS-CoV-2 spike-specific or RBD-specific IgG^+^ B cells. The antibody sequences and sequence features were shown in table S2. (**C**) Table shows the neutralization activities of three mAbs against VOCs and VOIs in pseudovirus neutralization assays. Experiments were done in 293T/ACE2 cells. The data are also shown in fig. S4, A and B. Data are representative of 2 biologically independent experiments for most VOCs and VOIs, but one experiment for BA.2 and BA.3. Each independent experiment contains 2 technical replicates. IC_50_ values were color-coded based on the key shown at the right. (**D**) Table shows the neutralization activities of three mAbs against VOCs in PRNT live virus neutralization assays. Data are representative of one independent experiment with 2 technical replicates. IC_50_ and IC_90_ values were color-coded based on the key shown at the right.

SARS-CoV-2 pseudovirus neutralization assays revealed that 3 of the RBD-binding mAbs potently neutralized the SARS-CoV-2 G614 virus including VHH7-5-82 (IC_50_: 38ng/ml), VHH7-7-53 (IC_50_: 68ng/ml), and SP1-77 (IC_50_: 20ng/ml) (Fig. 2C and fig. S4A). VHH7-5-82 also potently neutralized the Alpha (IC_50_: 20ng/ml) and Iota (IC_50_: 43ng/ml) VOCs (Fig. 2C and fig. S4A). VHH7-7-53 also potently neutralized Alpha (IC_50_: 43ng/ml), Epsilon (IC_50_: 62ng/ml), Iota (IC_50_: 43ng/ml) and Delta (IC_50_: 96 ng/ml) VOCs (Fig. 2C and fig. S4A). Notably, SP1-77 potently neutralized all currently known SARS-CoV-2 variants, including robust neutralization of the recently emerging Omicron sub-variants, BA.1 (IC_50_: 6.5 ng/ml), BA.2 (IC_50_: 33 ng/ml), BA.3 (IC_50_: 7 ng/ml), BA.4/BA.5 (IC_50_: 16 ng/ml) and BA.2.12.1 (IC_50_: 8 ng/ml) (Fig. 2C and fig. S4A). The IC_80_ neutralization values of SP1-77 against these variants were equally compelling (fig. S4B). Similar neutralization profiles for SP1-77 were found with an independent pseudovirus neutralization assay (fig. S4C). Robust neutralization of variants by SP1-77 also was seen in a plaque reduction neutralization test (PRNT) on live SARS-CoV-2 virus, with IC_50_ values ranging from 0.8 to 9.6 ng/ml (Fig. 2D). We also generated an SP1-77-derived antibody in which J_H_ and Jκ framework sequences outside of CDR3 were fully humanized and found that it retained similar robust neutralization activities against G614, Delta, Omicron sub-variants as SP1-77 (fig. S4D). Finally, we expressed the other four antibodies from the SP1 clonal lineage, each of which has unique pattern of somatic hypermutations compared to SP1-77. All have similar broad and potent neutralization activities (fig. S4D).

Surface plasmon resonance (SPR) confirmed interactions of the Fabs of the 3 RBD-binding nAbs with G614 S-2P SARS-CoV-2 spike protein (fig. S5A). Thus, we imaged Spike-Fab complexes using negative-stain electron microscopy (NSEM) to elucidate binding characteristics (fig. S5, B to D). Three-dimensional NSEM reconstructions indicated that VHH7-5-82, VHH7-7-53, and SP1-77 all bind to both up and down RBDs (fig. S5, B to E). However, these reconstructions further indicated that these 3 antibodies have distinct binding footprints. Thus, views of the RBD from the top, front, left, right and back showed that: VHH7-5-82 binds the top and right faces of RBD, overlapping with the RBM (fig. S6A); VHH7-7-53 binds the top face of the RBD, near the front, overlapping more widely with the RBM (fig. S6B); and SP1-77 binds the back and right faces, completely outside of the RBM (fig. S6C). Indeed, a surface plasmon resonance competition assay showed that SP1-77 does not compete with ACE2 for binding to the RBD or spike trimer (fig. S6D).

### cryo-EM structure of G614 S trimer in complex with the SP1-77 Fab

As SP1-77 neutralizes SARS-CoV-2 with remarkable potency and breadth, we determined cryo-EM structures of the full-length G614 S trimer in complex with the SP1-77 Fab (*29*) (fig. S7). 3D classification revealed two distinct conformations, including a three-RBD-down conformation and a one-RBD-up conformation, that both bound to three Fabs per S trimer (Fig. 3A) -- consistent with negative stain EM results (fig. S5D). These structures were refined to 2.7Å and 2.9Å resolution, respectively (fig. S7 to S9 and table S3). To improve resolution near the RBD/SP1-77 interface, we performed local refinement, leading to a 3.2 Å map that covers the Fab-RBD interface from the S trimer in the three-RBD-down conformation. From the S trimer in the one-RBD-up conformation, we obtained maps for the Fab-RBD interfaces in the down conformation and in the up conformation at 3.3 Å and 4.8Å resolution, respectively (fig. S7 to S9). These structures confirmed that the SP1-77 epitope sits on the opposite side of RBD from the ACE2 binding site (Fig. 3, B and C), consistent with the negative stain EM data and our surface plasmon resonance competition data showing that SP1-77 does not impact ACE2 binding.

**Figure 3.**
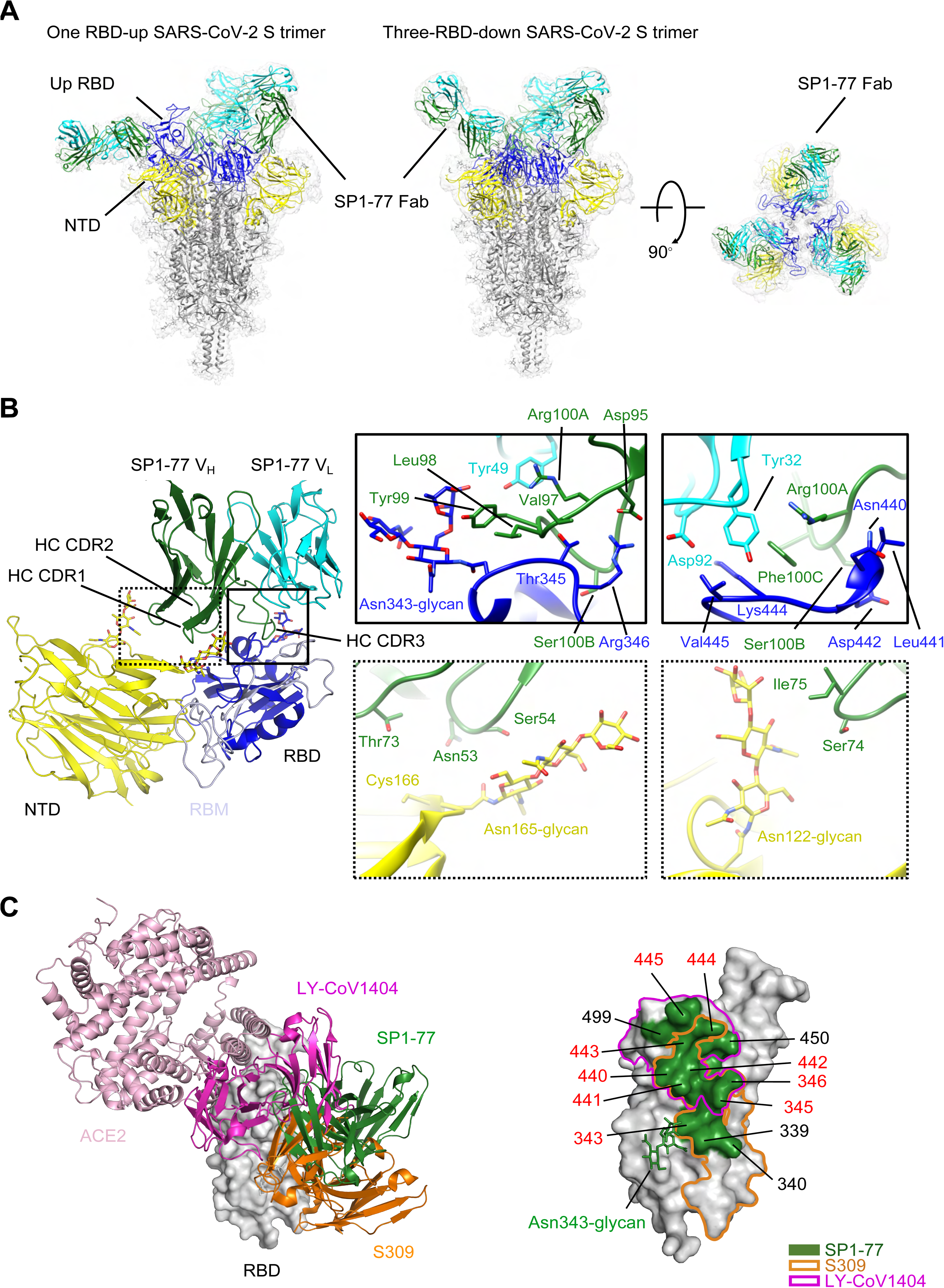
cryo-EM structures of SP1-77 Fab in complex of a full-length SASR-CoV-2 S trimer. (**A**) Cryo-EM structures of the SP1-77 Fabs in complex with the full-length S trimer in the one-RBD-up (2.9Å) and three-RBD-down (2.7Å) conformations, respectively. The EM density is colored in gray and the structures are represented in ribbon diagram with the RBD in blue, the NTD in yellow and the rest in dark gray. The heavy chain of SP1-77 is shown in green and the light chain is shown in cyan. There are three SP1-77 Fabs bound to one S trimer in both conformations. (**B**) Close-up view of the interactions between SP1-77 Fab and the RBD and NTD of the S trimer in the RBD-down conformation. Left, the HC CDR3 of SP1-77 makes the main contact with the RBD, with which away from receptor binding motif (RBM) colored in light blue, while the HC CDR2 touches the N-linked glycan from the NTD. Right, zoom-in views of the binding interface, showing the 17-residue HC CDR3 wedging into a groove formed by two segments of residues 339-346 and residues 440-445, respectively. The Asn343 glycan of the RBD also interacts with Tyr99 from HC CDR3. Two glycans at Asn122 and Asn165 in the NTD are in proximity to the HC CDR2 (Asn53, Ser54, Gly56, Thr57 and Asn58) and HC FW3 (Thr73, Ser74 and Ile75). (**C**) Comparison of binding mode and footprint of SP1-77 with ACE2 and other known neutralizing antibodies, including S309 and LY-CoV1404. Left, the RBD domain is in surface representation in gray. ACE2 and the antibodies in ribbon diagram with SP1-77 in green, S309 in orange, LY-CoV1404 in magenta and ACE2 in pink. Right, footprints on the RBD of various antibodies with the one for SP1-77 outlined in green, S309 in orange and LY-CoV1404 in magenta. The interface residues of the SP1-77 are indicated with the major contacting residues highlighted in red.

SP1-77 mainly makes direct contacts with the RBD, but also touches the N-terminal domain (NTD) from a neighboring protomer (Fig. 3B). The interaction buries an 939 Å^2^ surface area on the antibody and an 936 Å^2^ area from the spike protein. The HC and LC of SP1-77 contribute to 83% and 17% of the binding surface area, respectively. In particular, part of the 17-residue HC CDR3 wedges into a short groove formed by two segments in the RBD encompassing residues 343-346 and 441-444, respectively, while the tip of the 343-346 segment (mainly Thr345 and Arg346) projects into a ring-like structure formed by the HC CDR3 (Fig. 3B). The intertwined contacts between the HC CDR3 and the RBD account for 78% of the binding surface area contributed by the HC. There appears to be an extensive hydrogen bond network between Asn343, Thr345, Arg346, Asn440, Leu441 from the RBD and Val97, Leu98, Tyr99, Gly100, Arg100A, Ser100B from the HC CDR3. Notably, there is also a salt bridge between Arg346 of the RBD and Asp95 of the HC CDR3. Tyr99 of the HC CDR3 also packs against the fucose residue from the fucosylated Asn-343 glycan (*30*), probably forming a CH-π interaction, which buries a surface area of ~150 Å^2^ (Fig. 3B). In addition, there are a contacts between the 441-444 segment (Lys444 and Val445) and Tyr32 of the LC CDR1 and Asp92 of the LC CDR3 (Fig. 3B). Finally, the SP1-77 HC CDR2 and FW3 makes additional contacts with the Asn122 and Asn165 glycans on the NTD, probably by forming hydrogen bonds or water-mediated hydrogen bonds (Fig. 3B). Overall, the Cryo-EM structure revealed that, compared to other SARS-CoV-2 nAbs, SP1-77 binds a novel RBD epitope outside of the ACE2 binding motif via an interaction substantially mediated by the HC CDR3.

When the SP1-77 footprint is projected onto modeled RBDs from different SARS-CoV-2 variants (fig. S10, A and B), it is evident that most mutations are located outside of the SP1-77 binding site, except for the Arg346Lys in the Mu and Omicron-BA.1.1 variants (*31, 32*), and the Gly339Asp and Asn440Lys in all current Omicron sub-variants (*9, 33*). The modeled interfaces of SP1-77 bound to the Mu and Omicron RBDs, based on the SP1-77/G614 S complex, indicate that the conservative Arg346Lys mutation would preserve the interaction with HC CDR3 and not have much impact on binding. Likewise, the Gly339Asp and Asn440Lys mutations are at the edge of the SP1-77 epitope with their side chains pointing away from the binding interface, explaining why Omicron variants are sensitive to SP1-77 neutralization (fig. S10C). While mutations near the Asn343-glycan in Omicron reconfigure the orientation such that the carbohydrate projects away from the protein surface (*34*), such mutations have minimal impact on binding and neutralization activity of SP1-77 (fig. S10C). Finally, we also compared the SP1-77 footprint with the new mutations on the recently emerging Omicron sub-variants, BA.4, BA.5, BA.2.12 and BA.2.12.1. The R357K in BA.2.12 and F486V in BA.4/BA.5 are far from the SP1-77 footprint (fig. S10, A and B); while the L452Q mutation in BA.2.12.1 and L452R mutation in BA.4/BA.5 are close to but do not overlap with SP1-77 footprint. Thus, these structure studies reveal why SP1-77 potently and broadly neutralizes SARS-CoV-2 variants, including those that most recently emerged.

### SP1-77 has a novel neutralization mechanism

To address the SP1-77 neutralization mechanism, we applied a lattice light sheet microscopy (LLSM) approach to monitor target cell infection by a chimeric vesicular stomatitis virus in which the coat glycoprotein was replaced by the SARS-CoV-2 (Wuhan strain) S protein (VSV-SARS-CoV-2) (Fig.4 and fig. S11) (*35*). This chimeric virus is surface labeled with Atto565-NHS-ester (VSV-SARS-CoV-2-Atto 565) (fig. S11, A and B) and encodes a soluble eGFP that is expressed by infected cells. ACE2-expressing Vero cells that ectopically express TMPRSS2 protease (Vero TMPRSS2) were used as host cells. LLSM imaging revealed Atto 565-labeled viruses on cell surfaces at the earliest time points (Fig. 4, A to D, I, trajectories of viruses shown in magenta, light and dark blue). Single stepwise decreases in fluorescence intensity, corresponding to 25-30% of the total Atto 565 fluorescence, were observed for cell surface virus (Fig. 4, B to D, I, t2 to t3, magenta). As these step wise decreases are dependent on the presence of TPMRSS2 (*35*) and are also blocked by pharmacological inhibition of TMPRSS2 (Fig. 4J, central panel), they likely result from TMPRSS2 protease activity and, correspondingly, reflect S1 dissociation after S2’ cleavage. A subset of labeled chimeric viruses were internalized via endocytosis within a recorded 10-min time series (Fig. 4, A to D, I, trajectories of viruses shown in light and dark blue), followed by fluorescence-spreading, indicative of membrane fusion within the endosome (Fig. 4, B to D, I, t5 to t6, dark blue; see method). Under the conditions used in these experiments (*35*), spreading of Atto 565 was not observed on the cell surface.

**Figure 4.**
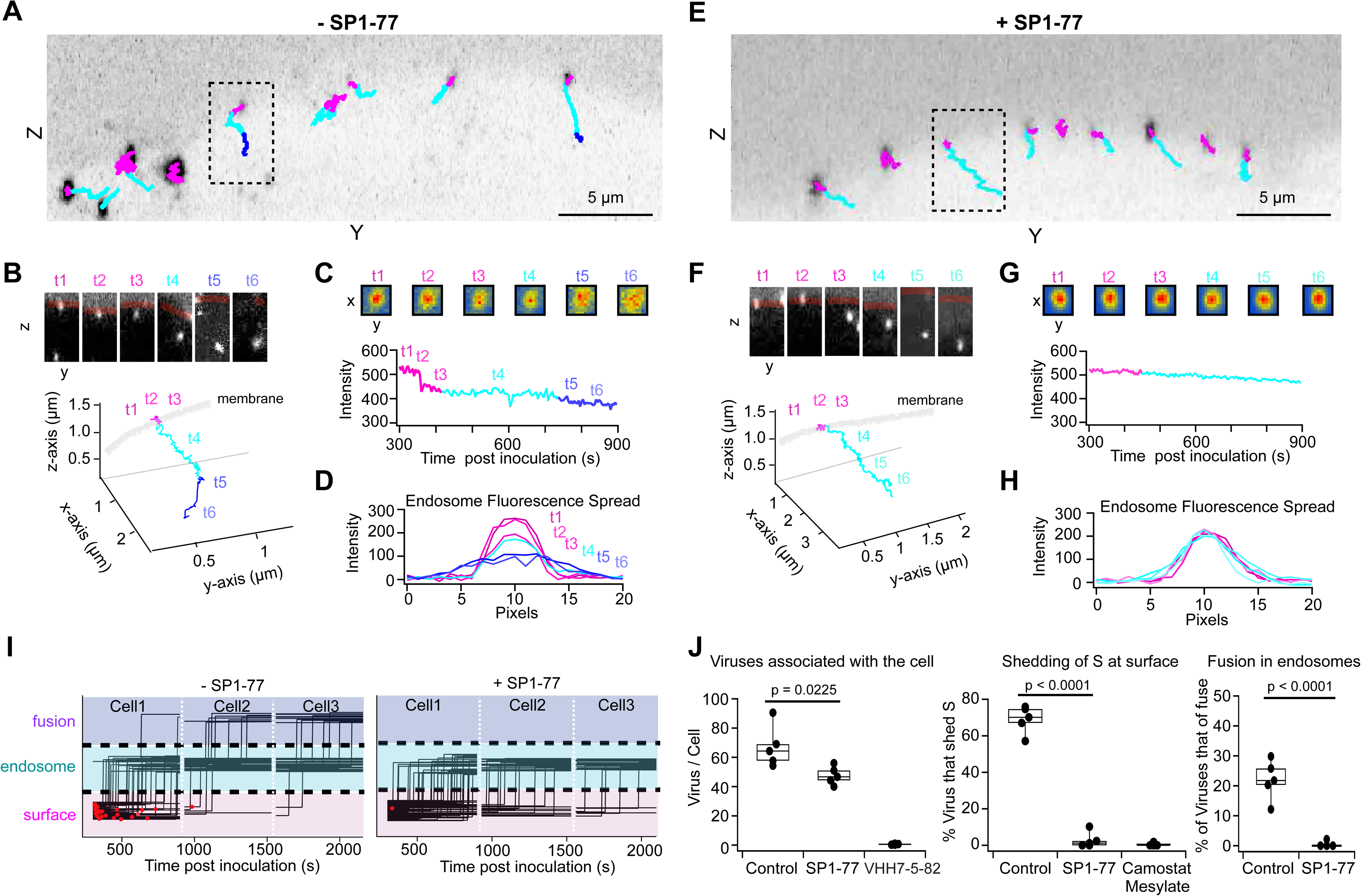
LLSM single virus tracking reveals SP1-77 inhibits S-fragment shedding and membrane fusion. (**A-I**) Trajectories of VSV-SARS-CoV-2-Atto 565 virus imaged every 4 sec with volumetric LLSM beginning 5 minutes after the start of a 4-min inoculation of Vero TMPRSS2 cells with virus at MOI ~ 2 without antibody treatment (**A-D**) or treated with 500 ng/mL SP1-77 (**E-H**). Trajectories color coded magenta when virus is localized to the cell surface, light blue when the virus internalizes into the cell, and dark blue if the Atto 565 signal is observed to spread from a point spread function (PSF) to a larger distribution indicating fusion of the viral envelope with an endosomal membrane. Single virus trajectories (**B, F**) with insets showing x-axis projection through 4 planes (top), 3D integrated intensity profiles shown in (**C, G**, bottom panel) and corresponding heat maps of fluorescence intensities from 4 plane z-projections in (**C, G**, top panel). Intensity line-profiles through the center of virions **(D, H**) appears as a single point spread function at the cell surface and in endosomes in the absence (**D**, t1-t4) of antibody or its presence (**H**, t1-t6) and which spreads out upon fusion of the virus envelope with endosomal membranes (**H**, t5-t6) in the control. (**I**) Summary of all single virus trajectories over the course of two single experiments, in the absence and presence of antibody, each from 3 cells imaged consecutively for 10 min. Red dots indicate virus trajectories where a TMPRSS2 dependent stepwise drop of Atto 565 signal was observed to occur which indicates release of a fragment of the S-protein. (**J**) Quantification of 5 experiments for each condition for the number of viruses in the total cell volume of every cell imaged (left), the number of Atto 565 drops observed in all the trajectories of virus when co-localizing to the cell surface (middle), and how many instances of Atto 565 dye spreading occurs within endosomes which indicated membrane fusion (right).

VSV-SARS-CoV-2-Atto 565 chimeric virus infectivity of Vero TMRSS2 cells was potently neutralized by its pre-incubation with SP1-77 IgG (IC_50_ ~15 ng/ml) (fig. S11E). Pre-incubation of VSV-SARS-CoV-2-Atto 565 with 500 ng/ml of SP1-77 did not appreciably impact its binding to the Vero TMPRSS2 cell surface (Fig. 4, E to I, trajectories of viruses shown in magenta and light blue). This finding is consistent with the SP1-77 footprint lying outside of the RBM and the inability of SP1-77 to block ACE2 engagement (Fig. 3C, fig. S6, C and D). In addition, pre-incubation with SP1-77 greatly diminished the number of Atto 565 fluorescence intensity step decreases of chimeric virus at the cell surface (Fig. 4, F to J, t2 to t3). This result indicates either the blockage of S2’ cleavage and/or inhibition of S1 dissociation. While pre-incubation of viruses with SP1-77 had modest impact on virus endocytosis (Fig. 4, F to I, trajectories of viruses shown in light blue), it dramatically inhibited fluorescence-spreading within the endosome --indicative of viral-endosomal membrane fusion block (Fig. 4, F to J). Together, these findings suggest that SP1-77 neutralizes the SARS-CoV-2 by blocking membrane fusion (Fig. 4, I and J). For comparison, LLSM single virion visualization revealed that pre-incubation with 500 ng/ml of our VHH7-5-82 antibody fully blocked binding of the chimeric virus to the cell surface (Fig. 4J), consistent with our findings that the VHH7-5-82 footprint overlaps with the RBM (fig. S6A).

To distinguish between the possible effect of SP1-77 on cleavage of S2’ versus dissociation of S1, we reconstructed these two processes *in vitro*. Activation of VSV-SARS-CoV-2-Atto 565 by brief treatment with trypsin for 30 minutes followed by the addition of the trypsin inhibitor aprotinin bypasses the requirement for host cell proteases TMPRSS2 and cathepsin L for infection, which indicates cleavage of the S2’ site by trypsin treatment (*35, 36*). Incubation of the trypsin-activated virus with recombinant ACE2 induces a signal decrease of ~75% of the total Atto565 fluorescence, as quantified by single particle imaging; while cleavage by trypsin alone or incubation with ACE2 alone does not induce this signal decrease (fig. S11, C and D). Assuming that an ~ 75% signal decrease *in vitro* reflects complete S1 dissociation from VSV-SARS-CoV-2 virus, then the corresponding 25-30 % signal decrease *in vivo* likely reflects S1 dissociation for a subset of spike proteins when the virus is attached to the cell surface (Fig. 4, B to D, t2 to t3). In contrast, the decrease in signal no longer occurred when trypsin-activated VSV-SARS-CoV-2-Atto 565 was incubated with SP1-77 and then exposed to ACE2 (fig. S11, C and D). This finding indicates that SP1-77 inhibits ACE2-induced dissociation of S1 from the pre-cleaved S1/S2 complex. Finally, the SP1-77 Fab also inhibits S1 dissociation in this assay and, corresponding, robustly neutralizes VSV-SARS-CoV-2 virus (fig. S11, D and E). This finding indicates that SP1-77 can achieve it neutralization activity by blocking S1 dissociation via a monovalent interaction. See fig. S12 for a model of the SP1-77 neutralization mechanism.

## DISCUSSION

The RBD epitope targeted by SP1-77 overlaps very modestly with that of the LY-CoV1404 (Bebtelovimab) SARS-CoV-2 neutralizing nAb that is currently approved for therapeutic use. The RBD-binding site of LY-CoV1404 includes the 440-445 segment of the RBD that is also bound by SP1-77 (*37*) (Fig. 3C). Like SP1-77, LY-CoV1404 potently neutralizes all currently known VOCs. However, the bulk of the LY-CoV1404 footprint overlaps with the RBM, which mechanistically allows LY-CoV1404 to achieve neutralization by blocking ACE2 binding (*37*). Like SP1-77, the S309 nAb and its Sotrovimab derivative previously used therapeutically bind outside of the RBM (*17*). However, S309 is quite distinct from SP1-77 in that it fails to neutralize Omicron BA.2 and is less potent than SP1-77 in neutralizing other VOCs (*38*). In this regard, the S309 footprint covers both the 339-346 and 440-451 segments that are also in the SP1-77 footprint, with the long axis of the S309 footprint aligning along the exposed ridge of the RBD in the closed S trimer (Fig. 3C). Notably, however, the same axis of the SP1-77 footprint is almost perpendicular to that of S309, which makes it possible for SP1-77 to reach the NTD (Fig. 3B). Indeed, SP1-77 has a unique binding mode, not seen among any of the previously characterized SARS-CoV-2 nAbs, that may underlie its robust and broadly potent neutralization activity.

The SP1-77 neutralization mechanism we describe provides the first insight into how a non-ACE2 blocking antibody can potently neutralize SARS-CoV-2. Besides S309, several other Omicron sub-variant nAbs that have binding epitopes outside of the RBM have been reported very recently (*8, 39*). However, like S309, the mechanism by which these antibodies neutralize SARS-CoV-2 has not been described. The LLSM-based approach we employed to elucidate the novel SP1-77 mechanism should be readily applicable for mechanistic studies of other neutralizing antibodies that do not function via an ACE2 blocking mechanism. The unique binding characteristics and novel neutralization mechanism of the SP1-77, coupled with its broad and potent neutralization of SARS-CoV-2 variants, suggests that it could be a therapeutic against current and, very likely, newly-arising VOCs. Also, SP1-77 might be useful in a cocktail with other nAbs, such as LY-CoV1404, that potently neutralize all tested VOCs through an ACE2-blocking mechanism. In addition, the novel neutralization mechanism of SP1-77 may inspire the design of new vaccine strategies to more broadly target BA.2 and other VOCs.

The use of a solely human V_H_1-2-rearranging mouse to obtain SP1-77 posed a rigorous test for this class of mouse model. In this regard, all 9 well-characterized V_H_1-2-based SARS-CoV-2 neutralizing antibodies isolated from prototype SARS-CoV-2 virus-infected human patients bind a similar epitope on the spike protein RBD domain (*15, 16, 22, 40–44*) (fig. S13, A and B). None of these antibodies tested neutralized Beta, Gamma or Omicron variants, due to their sensitivity to the E484K/A mutation in these variants (*15, 16, 22, 40–44*). Indeed, the striking binding and neutralization similarities of these 9 V_H_1-2-based nAbs suggested that most V_H_1-2-based SARS-CoV-2-neutralizing antibodies in infected humans are selected by a common prototype neutralization pathway mediated by predisposition of the germline-encoded V_H_1-2 variable region to bind to the RBD (*22*). Notably, the footprints of our 3 V_H_1-2-based SARS-CoV-2 nAbs are distinct from those of the 9 well-characterized V_H_1-2 based SARS-CoV-2 nAbs isolated from human patients. Thus, our solely V_H_1-2-rearranging mouse model permits isolation of human V_H_1-2-based SARS-CoV-2 nAbs with novel binding characteristics relative to those derived from patients, potentially by placing greater emphasis on the contribution of CDR3 with respect to diversification of the primary BCR repertoire.

The V_H_1-2/Vκ1-33-rearranging mouse model has served as a very successful prototype for documenting the potential utility of this humanized antibody-discovery approach by providing the broad, potent, and mechanistically-distinct SP1-77 nAb. Going forward, this type of mouse model could be readily extended to other human V_H_ and V_L_ segments, to augment its potential beyond that which we demonstrate in this striking initial study. Finally, this new approach for induction of humanized antibodies from mice could help in the generation of novel therapeutic antibodies and vaccine strategies against other diverse pathogens for which rare cross-reactive neutralizing antibodies are needed such as HIV-1 and influenza.

## Author contributions

S.L. and F.W.A. initiated and oversaw this overall study. S.L., T.K., A.K. B.C., B.F.H. and F.W.A. designed the experiments. S.L. and H.D. generated the mouse model. S.L., C.J., and M.T. performed immunizations, generated the antibodies, and characterized the antibodies. K.S. provided the spike immunogen and N.P. and H.P. provided the VHH7-RBD immunogen. R.P., M.B. and S.L. performed binding experiments. D.M., A.E., S.K. and L.B. performed the pseudovirus neutralization experiments. G.S. performed the PRNT neutralization experiments. M.A. and K.C. performed the SPR binding and competition analyses. R.E. and K.M. performed the NSEM experiments and interpreted that data. J.Z., Q.P. and B.C. performed the Cryo-EM experiments and interpreted the data. A.K., A.S. and T.K. performed the live imaging experiment with lattice light sheet microscopy and interpreted their findings in the context of the neutralization mechanism. A.Y.Y. performed the CDR3 diversity analyses. A.C.W. performed ES cell injections. L.V.F. collected blood samples and performed mouse maintenance. S.L., J.Z., A.K. T.K., B.C., B.F.H. and F.W.A. designed figures and wrote the manuscript. All other authors including helped polish the manuscript.

## Data and materials availability

All newly created data and materials described in this manuscript will be available from the corresponding author upon reasonable request. Nucleotide sequences have been deposited to GenBank (accession Nos. ON568518 to ON568583). NSEM reconstructions generated during this study are available at EMD-27043, EMD-27044 and EMD-27046. Cryo-EM reconstructions and atomic models generated during this study are available at the PDB and EMBD databases under the following accession codes: 7UPW,7UPX,7UPY, EMD-26676 (three RBD-down D614G spike in complex with SP1-77 Fabs), EMD-26677 (local refinement of RBD in complex with SP1-77 fab), EMD-26678 (one RBD-up D614G spike in complex with SP1-77 Fabs). The next-generation sequencing data reported in this paper have been deposited in the Gene Expression Omnibus (GEO) database under GSE197255. This work is licensed under a Creative Commons Attribution 4.0 International (CC BY 4.0) license, which permits unrestricted use, distribution, and reproduction in any medium, provided the original work is properly cited. To view a copy of this license, visit https://creativecommons.org/licenses/by/4.0/. This license does not apply to figures/photos/artwork or other content included in the article that is credited to a third party; obtain authorization from the rights holder before using such material.

## Materials and Methods

### Mice and Embryonic Stem Cells

All mouse experiments were performed under protocol 20-08-4242R approved by the Institutional Animal Care and Use Committee of Boston Children’s Hospital. Mice were maintained on a 14-h light, 10-h dark schedule in a temperature-controlled environment, with food and water provided ad libitum. ES cells were grown on a monolayer of mitotically inactivated mouse embryonic fibroblasts (iMEF) in DMEM medium supplemented with 15% bovine serum, 20mM HEPES, 1x MEM nonessential amino acids, 2mM Glutamine, 100 units of Penicillin/Streptomycin, 100 mM ß-mercaptoethanol, 500 units/ml Leukemia Inhibitory Factor (LIF).

### Generation and Characterization of the V_H_1-2/Vκ1-33-rearranging mouse model

All the genetic modifications were introduced into previously generated V_H_1-2^IGCR1Δ^ ES cells (129/Sv and C57BL/6 F1 hybrid background) (*1*), using targeting strategies described previously (*1, 2*). In the *Igh* locus, the deletion of all mouse V_H_s was mediated by two guide RNAs that, respectively, target sequences ~100 bp upstream of the most distal mouse V_H_1-86P and ~4.3 kb upstream of the human V_H_1-2 that replaced V_H_5-2 in this modified ES cell line, respectively. In the *Igκ* locus, the mouse Vκ3-2 segment was replaced with human Vκ1-33 segment with an attached CTCF-binding element (CBE) (<u>atccaggaccagcagggggcgcggagagcaca</u> <u>ca</u>) inserted 50 bp downstream of human Vκ1-33 segment. The Vκ1-33/CBE replacement of Vκ3-2 was mediated by homologous recombination using a PGKneolox2DTA.2 (Addgene #13449) construct and two guide RNAs that target the mouse Vκ3-2 segment. The Cer/sis regulatory element in this ES cell line was deleted via the use of two guide RNAs that target sequences ~60 bp upstream and ~1 kb downstream of Cer/sis. A human TdT gene is expressed in this mouse model. The human TdT cDNA was cloned into CTV (Addgene #15912) construct in which the TdT expression was driven by CAG promotor and followed by a EGFP expression that mediated by an internal ribosome entry site (IRES) (*3*). The TdT expression cassette was inserted into the first intron of mouse Rosa26 gene which is on the same chromosome 6 with *Igκ* locus by homologous recombination. The sequences of guide RNAs used for targeting are listed in table S4. The modifications on the chromosome 12 (the *Igh* loci) and chromosome 6 (the *Igκ* and Rosa26 loci) were made independently in two ES clones. We performed blastocyst injection of these two ES clones, respectively, generating the V_H_1-2^mVHΔ/IGCR1Δ^ - rearranging mouse model and the V_H_1-2^IGCR1Δ^/Vκ1-33^CSΔ^ - rearranging mouse model. The V_H_1-2^mVHΔ/IGCR1Δ^/Vκ1-33^CSΔ^ - rearranging F1 mice were generated by cross-breeding of these two colonies. We then generated mice that were homozygous for each modification and used these mice for all experiments.

To characterize B cells in the homozygous V_H_1-2^mVHΔ/IGCR1Δ^/Vκ1-33^CSΔ^ - rearranging mouse model, splenocytes were isolated from 5-8 week old mice and stained with the following antibodies: APC anti-B220 (eBioscience 17-0452-83); PE anti-Thy1.2 (eBioscience 12-0902-83); Bv605 anti-IgM (BioLegend 406523); Bv711 anti-IgD (BioLegend 405731).

### Splenic B Cell Purification and HTGTS-Rep-seq Analysis

Splenic B cells were purified from 5-8 week old mice by MACS^®^ Microbeads according to the manufacturer’s protocol. In brief, spleens were dissected out from unimmunized mice, prepared into single cell suspensions and incubated with anti-B220 Microbeads for 20 minutes at 4°C. Then, the splenic B cells were collected using the LS column and MACS^TM^ Separater and lysed for genomic DNA extraction. 10 ug of genomic DNA from purified splenic B cells was used for generating HTGTS-rep-seq libraries as previously described (*4*). Four bait primers that target mouse J_H_1, J_H_2, J_H_3 and J_H_4 were mixed to capture the *Igh* heavy chain repertoire in one library. Likewise, four 4 bait primers that target mouse Jκ1, Jκ2, Jκ4 and Jκ5 were mixed to capture the *Igκ* light chain repertoire in one library. The sequences of mouse J_H_ and Jκ primers were as same as the previously reported (*5*). These HTGTS-rep-seq libraries were sequenced by Illumina NextSeq 2 x 150-bp paired end kit and analyzed with the HTGTS-Rep-seq pipeline (*6*).

### ELISA

96 well plates (Thermo Fisher 446612) were coated with SARS-CoV-2 spike, RBD, NTD or SARS-CoV-1 spike at 100 ng/well at 4°C overnight, and blocked with blocking buffer (150 mM NaCl, 10 mM NaPO_4_ pH 7.2, 1% bovine serum albumin) at room temperature for one hour. Then, the plates were washed with the washing buffer (150 mM NaCl, 10 mM NaPO_4_ pH 7.2, 0.05% Tween 20) to remove the blocking buffer. Sera were serial diluted from 1:200 in the blocking buffer and added to the plates for 1-hour incubation at room temperature. The plates were washed again with washing buffer. Next, 1:1000 diluted AP-conjugated anti-mouse IgG antibody (Southern Biotech, 1030-04) (Pooled sera: reacts with the heavy chains of mouse IgG1, IgG2a, IgG2b, IgG2c, and IgG3) was added and plates were incubated for 1-hour at room temperature. The plates were washed again and phosphatase substrate (Sigma S0942) was added to each well. The absorbance at 405nm (OD405) was recorded. Background absorbance was determined based on wells blotted with the blocking buffer.

### Immunogens and Immunizations

HexaPro Spike immunogen was produced and purified as previously described (*7–10*). HexaPro Spike was stabilized by the introduction of six prolines in the S2 region and contained an HRV 3C-cleavable C-terminal twinStrepTagII-8×His tag to facilitate purification (PMID: 32703906). DNA encoding SARS-CoV-2 HexaPro Spike was synthesized (Genscript) and transiently transfected in FreeStyle 293F cells (Thermo Fisher) using 293Fectin (ThermoFisher). On day 5, cell culture supernatants were harvested by centrifugation of the culture followed by filtering through a 0.8-µm filter. Supernatant was concentrated down to 400 mL and Spike protein was purified by StrepTactin chromatography (IBA) using the manufacturer’s buffers. Trimeric spike protein was further purified by Superose 6 size-exclusion chromatography (GE Healthcare) in 10 mM Tris pH 8, 500 mM NaCl. All purification steps were done at room temperature. Purified proteins were quantified and snap frozen in dry ice mixed with ethanol.

The VHH7-RBD immunogen was made as previously described (*11*). Briefly, VHH7-RBD DNA was synthesized (Integrated DNA Technologies) and assembled in the pVRC vector. Constructs were transfected into Expi293 cells (Thermo Fischer Scientific) using Polyethyleneimine “Max” (Polysciences). Cell cultures were maintained in Expi293 Media (Thermo Fischer Scientific) at 37°C for 4 day following transfection. Proteins were harvested by centrifugation at 5,000 Å~ g for 30 min at 4°C, followed by affinity chromatography with HisPur Ni-NTA Resin (Thermo Fischer Scientific) and size exclusion chromatography with a Hi-Load 16/600 S75 column (Cytivia).

For each immunization, 200 ul of immunogen mix, containing 25 ug of filter-sterilized protein immunogen and 60 ug of Poly I:C in PBS, was injected into the peritoneum of each mouse. 8 weeks old mice were immunized twice with same immunogen, 4 weeks apart. Blood samples were collected one day before the first immunization and two weeks after each immunization. Spleens were collected 3 weeks after the second immunization.

### Antigen-specific B Cell Sorting and Single Cell RT-PCR

Mouse spleen samples from immunized mice were processed for single B cell sorting as previously described (*5*). In brief, single cell suspension of splenocytes were stained with several fluorescent antibodies. Antigen-specific IgG^+^ B cells were selected for the phenotype B220^+^ (PE-Cy7: BD 552772), IgD^-^ (BV711: BioLegend 103255), IgG1^+^ (BV605: BD 563285), IgG2a^+^ (BV605: BD 564024), Spike^+^ (Alex647) or RBD^+^ (Alex647). SARS-CoV2 spike and RBD were conjugated with Alex647 fluorescence (Thermo Fisher A30009).

Single cell RT-PCR was performed as described previously (*1*). In brief, single antigen-specific IgG^+^ B cells were sorted into 96-well plate that contain 5ul of lysis buffer in each well. After sorting, we used a primer mixture that specifically targets Cu, Cγ1, Cγ2a and Cκ to perform reverse transcription and then two rounds of nested PCR to amplify the V(D)J sequences of rearranged V_H_1-2 and Vκ1-33 exons. PCR products were run on agarose gels and subjected to Sanger sequencing to confirm their identity. Primer sequences used are listed in Supplementary table S4.

### Monoclonal Antibody and Fab Production

Sequences of paired V_H_1-2-based and Vκ1-33 based variable region exons of antigen-specific antibodies were determined by sanger sequencing following single cell RT-PCR (table S2). The DNA fragments containing the heavy-chain and the light-chain variable region exons (table S2), with human constant region sequences (IgG4 or IgG1, Igκ) at the C terminus were synthesized in Integrated DNA Technologies (gblock) and cloned into pcDNA3.4^+^ plasmids, respectively. Monoclonal antibodies were generated using the Expi293 expression system (Thermo Fisher Scientific) according to the product manual. The antibodies were purified from Expi293 cell supernatant by high-performance liquid chromatography (HPLC) coupled with HiTrap Protein A HP columns (Cytiva, formerly GE Healthcare). Monoclonal antibodies were then dialyzed by SnakeSkin dialysis tubing (10K MWCO, 22 mm) and stored at 4 °C in PBS.

Fabs were made as previously described (*12*). Full-length antibodies were dialyzed into 20 mM sodium phosphate, 10 mM EDTA, pH 7.0 and then concentrated to ~20 mg/ml by centrifugal filtration (Amicon/EMD Millipore). IgG antibodies were digested in 20 mM sodium phosphate, 10 mM EDTA, 20 mM cysteine, pH 7.4 with papain-agarose resin (Thermo Fisher Scientific) for 5 hr at 37°C. The Fab fragments are then separated from non-digested IgG and Fc fragments by a 1 hr incubation with rProtein A Sepharose Fast Flow (GE Healthcare). Post incubation, cystein was removed by centrifugal filtration and buffer exchanged into 1x PBS pH7.4 (Amicon/EMD Millipore).

### Plaque Reduction Neutralization Test (PRNT)

SARS-CoV-2 Plaque Reduction Neutralization Test (PRNT) were performed in the Duke Regional Biocontaiment Laboratory BSL3 (Durham, NC) as previously described with virus-specific modifications (*13*). Briefly, two-fold dilutions of a test sample (e.g. serum, plasma, purified Ab) were incubated with 50 PFU SARS-CoV-2 virus (Isolate USA-WA1/2020, NR-52281) for 1 hour. The antibody/virus mixture is used to inoculate Vero E6 cells in a standard plaque assay (*14, 15*). Briefly, infected cultures are incubated at 37°C, 5% CO2 for 1 hour. At the end of the incubation, 1 mL of a viscous overlay (1:1 2X DMEM and 1.2% methylcellulose) is added to each well. Plates are incubated for 4 days. After fixation, staining and washing, plates are dried and plaques from each dilution of each sample are counted. Data are reported as the concentration at which 50% of input virus is neutralized. A known neutralizing control antibody is included in each batch run (Clone D001; SINO, CAT# 40150-D001).

### Antibody Affinity and Competition Assays by Surface Plasmon Resonance (SPR)

SPR screening and affinity measurements of monoclonal Fabs binding to SARS-CoV-2 spike proteins were performed using a Biacore S200 instrument (Cytiva, formerly GE Healthcare) in HBS-EP+ 1X running buffer. The SARS-CoV-2 spike (S) proteins with a Strep tag were first captured onto a Streptavidin sensor chip to a level of 200-400 RU. For screenings, the Fabs were injected over the captured S proteins using the high performance injection mode at a flow rate of 30uL/min. Association phase was maintained at 180s injections of each Fab followed by a dissociation of 720s. After each dissociation phase, the S proteins and bound Fabs were removed from the sensor surface using a 30s regeneration pulse of Glycine pH1.5. For affinity measurement, the Fabs were injected over the captured S proteins using single cycle kinetics with the high performance injection mode at a flow rate of 50uL/min. Association phase was maintained at 120s injections of each Fab at increasing concentrations followed by a dissociation of 900s. After each dissociation phase, the CoV-S proteins and bound Fabs were removed from the sensor surface using a 30s regeneration pulse of Glycine pH1.5. Results were analyzed using the Biacore S200 Evaluation software (Cytiva). A blank streptavidin surface along with blank buffer binding were used for double reference subtraction to account for non-specific protein binding and signal drift. Subsequent curve fitting analyses were performed using a 1:1 Langmuir model with a local Rmax for the Fabs or using the heterogeneous ligand model with local Rmax.

Antibody binding competition and blocking were measured by SPR following immobilization by amine coupling of monoclonal antibodies to CM5 and CM3 sensor chips (BIAcore/Cytiva) (*7*). Antibody blocking assays were performed by sequential high performance injections of Spike protein (20 µM) over mAb immobilized surfaces for 3 minutes at 30 µL/min immediately followed by a test Ab (200 µM) for 3 minutes at 30 µL/min. The dissociation of the antibody sandwich complex with the spike protein was monitored for 10 minutes with buffer flow and then a 24 second injection of Glycine pH2.0 for regeneration. Double reference subtraction was used to account for signal drift. Data analyses were performed with BIA-evaluation 4.1 software (BIAcore/Cytiva). The reported competition experiments are representative of two data sets.

### Pseudovirus Neutralization Tests

The pseudovirus neutralization assay in 293T/ACE2 cells performed at Duke has been described in detail and is a formally validated adaptation of the assay utilized by the Vaccine Research Center; the Duke assay is FDA approved for D614G. For measurements of neutralization, pseudovirus was incubated with 8 serial 5-fold dilutions of antibody samples (1:20 starting dilution using antibodies diluted to 1.0 mg/ml or 0.5 mg/ml) in duplicate in a total volume of 150 µl for 1 hr at 37°C in 96-well flat-bottom culture plates. 293T/ACE2-MF cells were detached from T75 culture flasks using TrypLE Select Enzyme solution, suspended in growth medium (100,000 cells/ml) and immediately added to all wells (10,000 cells in 100 µL of growth medium per well). One set of 8 wells received cells + virus (virus control) and another set of 8 wells received cells only (background control). After 71-73 hrs of incubation, medium was removed by gentle aspiration and 30 µl of Promega 1X lysis buffer was added to all wells. After a 10-minute incubation at room temperature, 100 µl of Bright-Glo luciferase reagent was added to all wells. After 1-2 minutes, 110 µl of the cell lysate was transferred to a black/white plate. Luminescence was measured using a GloMax Navigator luminometer (Promega). Neutralization titers are the inhibitory dilution (ID) of serum samples at which RLUs were reduced by 50% (ID50) compared to virus control wells after subtraction of background RLUs. Serum samples were heat-inactivated for 30 minutes at 56°C prior to assay.

The pseudovirus neutralization assays in 293T/ACE2/TMPRSS2 cells were performed as previously described (*16–18*). Briefly, human codon-optimized cDNA encoding SARS-CoV-2 spike glycoprotein of the WA-1/2020 and variants were synthesized by GenScript and cloned into eukaryotic cell expression vector pcDNA 3.1 between the *BamH*I and *Xho*I sites. Pseudovirions were produced by co-transfection Lenti-X 293T cells with psPAX2(gag/pol), pTrip-luc lentiviral vector and pcDNA 3.1 SARS-CoV-2-spike-deltaC19, using Lipofectamine 3000. The supernatants were harvested at 48h post transfection and filtered through 0.45µm membranes and titrated using 293T/ACE2/TMPRSS2 cells (HEK 293T cells that express ACE2 and TMPRSS2 proteins). For the neutralization assay, 50 µL of SARS-CoV-2 S pseudovirions (counting ~200,000 relative light units) were pre-incubated with an equal volume of medium containing serial dilutions of mAbs at room temperature for 1h. Then 50 µL of virus-antibody mixtures were added to 293T-ACE2-TMPRSS2 cells (10^4^ cells/50 µL) in a 96-well plate. The input virus with all SARS-CoV-2 strains used in the current study were the same (2 x 10^5^ relative light units/50 µL/well). After a 3 h incubation, fresh medium was added to the wells. Cells were lysed 24 h later, and luciferase activity was measured using One-Glo luciferase assay system (Promega, Cat# E6130). The assay of each serum was performed in duplicate, and the 50% neutralization titer was calculated using Prism 9 (GraphPad Software). Controls included cells only, virus without any antibody and positive sera.

### NSEM Methods

To form the Fab-spike complex, Fab was mixed with spike at a 9:1 molar ratio and incubated at 37 °C for 1 h. The complex was then cross-linked for 5 min by diluting to 200 µg/ml spike concentration with 8 mM glutaraldehyde in HBS buffer (20 mM HEPES, 150 mM NaCl, 5 g/dL glycerol, pH 7.4), and then quenched for 5 min by addition of sufficient 1 M Tris stock to give 80 mM final Tris concentration. For negative stain, a portion of the sample was diluted in HBS buffer to 100 or 50 µg/ml and 5 µl applied to a glow-discharged carbon-coated EM grid for 10-12 second, then blotted, and stained with 2 g/dL uranyl formate for 1 min, and then blotted and air-dried. Grids were examined on a Philips EM420 electron microscope operating at 120 kV and nominal magnification of 49,000x, and ~100 images were collected on a 76 Mpix CCD camera at 2.4 Å/pixel. Images were analyzed by 2D class averages and 3D reconstructions calculated using standard protocols with Relion 3.0 (*19*).

### Cryo-EM Sample Preparation and Data Collection

Cryo-EM images were acquired on a Titan Krios electron microscope equipped with a Gatan K3 direct electron detector. We used cryo-SPARC (*20*) for particle picking, two-dimensional (2D) classification, three dimensional (3D) classification and refinement. To prepare cryo EM grids, the full-length G614 spike trimer (3.1 mg/ml)(*21*) and the SP1-77 Fab (7 mg/ml) were mixed at a molar ratio of 1:9 and incubated at room temperature for one hour. The complex was purified by gel filtration chromatography on a Superose 6 10/300 column (GE Healthcare, Chicago, IL) in a buffer containing 25 mM Tris-HCl, pH 7.5, 150 mM NaCl, 0.02% DDM. Both the mixed complex and purified complex were used for cryo grid preparation. 4.0 µl of the complex was applied to a 1.2/1.3 Quantifoil gold grid (Quantifoil Micro Tools GmbH), which had been glow discharged with a PELCO easiGlow^TM^ Glow Discharge Cleaning system (Ted Pella, Inc.) for 60 s at 15 mA. Grids were immediately plunge-frozen in liquid ethane using a Vitrobot Mark IV (ThermoFisher Scientific), and excess protein was blotted away by using grade 595 filter paper (Ted Pella, Inc.) with a blotting time of 4 s, a blotting force of −12 at 4°C with 100% humidity. The grids were first screened for ice thickness and particle distribution. Selected grids were used to acquire images by a Titan Krios transmission electron microscope (ThermoFisher Scientific) operated at 300 keV and equipped with a BioQuantum GIF/K3 direct electron detector. Automated data collection was carried out using SerialEM version 3.8.6 (*22*) at a nominal magnification of 105,000× and the K3 detector in counting mode (calibrated pixel size, 0.83 Å) at an exposure rate of 13.761 electrons per pixel per second. Each movie add a total accumulated electron exposure of ~53.853 e-/Å^2^, fractionated in 50 frames. Data sets were acquired using a defocus range of 0.5-2.2 µm.

### Image Processing and 3D Reconstructions

Cryo-EM images were first processed in cryoSPARC v.3.3.1.(*20*) with the drift correction performed using the patch mode, and contrast transfer function (CTF) estimated by the patch mode as well. Motion corrected sums with dose-weighting were used for subsequent image processing. Templates for particle-picking were generated from a small number of manually picked particles and the template-based particle picking was then performed for all 28,899 recorded images with 7,657,522 particles extracted in total (box size 672Å, downsizing to 128Å). The particles were subjected to four rounds of 2D classification in cryoSPARC, giving 3,894,421 good particles. A low-resolution negative-stain reconstruction of the Wuhan-Hu-1 (D614) sample was low-pass filtered to 40Å resolution and used as an initial model for 3D classification (*23*). The selected good particles were subjected to two rounds of heterogeneous 3D classification with six copies of the initial model as references in C1 symmetry. One major class (12.8%) with clear structural features was re-extracted to a smaller boxsize (480Å) and subjected to another two rounds of heterogeneous refinement with four copies of the initial model as references in C1 symmetry. There were two major classes, one representing the three-RBD-down S trimer bound with three SP1-77 Fabs and another representing the one-RBD-up S trimer in complex with three SP1-77 Fabs. The first class was subjected to one round of non-uniform refinement in C3 symmetry, giving a map at 2.9Å resolution from 250,319 particles. The second class was refined in C1 symmetry instead, giving a map at 3.1Å resolution from 209,249 particles. To further improve the overall resolution, both classes were subjected to local CTF refinement, resulting in a 2.7Å map for the complex with the S trimer in the three-RBD-down conformation, and a 2.9Å map for a 2.7Å map for the complex with the S trimer in the one-RBD-up conformation. To improve the density for the Fab and RBD, several different local refinements were performed with a soft mask covering a single RBD and the bound Fab. From the three-RBD-down complex, the local refinement gave a map at 3.2Å resolution; from the one-RBD-up complex, the local refinements gave a 3.3Å map for the Fab bound to the RBD in the down conformation, and a 4.8Å map for the Fab bound to the RBD in the up conformation. The best density maps were used for model building.

All resolutions were reported from the gold-standard Fourier shell correlation (FSC) using the 0.143 criterion. Density maps were corrected from the modulation transfer function of the K3 detector and sharpened by applying a temperature factor that was estimated using Sharpening Tools in cryoSPARC. Local resolution was also determined using cryoSPARC.

### Model Building

For model building, we used our G614 S trimer structures (PDB ID: 7KRQ and PDB ID: 7KRR (*24*)) as the initial templates for the spike protein, and the S309 antibody structure (PDB ID: 6WS6 (*25*)) for SP1-77 antibody. We predicted the SP1-77 Fab structure by AlphaFold (*26*), largely confirming our model building and the predicted structure also helped further improving some details of the final model. Several rounds of manual building were performed in Coot (*27*). The models were refined in Phenix (*28*) against the 2.7Å (three-RBD-down) and 2.9Å (one-RBD-up) maps, respectively, as well as the 3.2Å map from the masked local refinement using the model containing a single RBD and the SP1-77 Fab. Iteratively, refinement was performed in both Phenix (real space refinement) and ISOLDE (*29*), and the Phenix refinement strategy included minimization_global, local_grid_search, and adp, with rotamer, Ramachandran, and reference-model restraints, using 7KRQ and 7KRR as the reference model. The refinement statistics are summarized in table S3. Structural biology applications used in this project were compiled and configured by SBGrid (*30*).

### Generation and purification of VSV-SARS-CoV-2 chimeras

The generation of recombinant VSV chimeras expressing eGFP where the glycoprotein G was replaced with spike (S) protein Wuhan-Hu-1 strain (VSV-SARS-CoV-2) was previously described (*31*). VSV-SARS-CoV-2 chimera were obtained by infection of MA104 cells. Cells were grown in 15 to 20 150-mm dishes and infected at a multiplicity of infection (MOI) of 0.01. The supernatant containing the virus was collected 72 hours post infection and clarified by centrifugation at 1,000 x g for 10 min at 4°C. A pellet with virus and extracellular particles was obtained by centrifugation in a Ti45 fixed-angle rotor at 72,000 x g (25,000 rpm) for 2 hours at 4°C, then resuspended overnight in 0.5 mL PBS at 4°C. This solution was layered on top of then pelleted through 15% sucrose-PBS solution by centrifugation in a SW55 swinging-bucket rotor at 148,000 x g (35,000 rpm) for 2 hours at 4°C. The pellet was then resuspended overnight in 0.4 mL PBS at 4°C, layered on top of a 15 to 45% sucrose-PBS linear gradient and subjected to centrifugation in a SW55 swinging-bucket rotor at 194,000 x g (40,000 rpm) for 1.5 hours at 4°C. The predominant light scattering band located in the lower one-third of the gradient and containing the virions was removed by side puncture of the gradient tube. Approximately 0.3 ml of this solution was mixed with 25 ml of PBS and subjected to centrifugation in a Ti60 fixed-angle rotor at 161,000 x g (40,000 rpm) for 2 hours at 4°C. The final pellet was resuspended overnight in 0.2 - 0.5 mL PBS aliquots and stored at −80°C.

### VSV-SARS-CoV-2 Atto 565 labeling

Stock concentration of VSV-SARS-CoV-2 at a concentration of ~150 µg/mL viral RNA was conjugated with Atto565-NHS ester (Sigma-Aldrich, cat.72464) as previously described (*32, 33*). Virus stocks in 200 µL were adjusted to 0.1 M NaHCO3 (pH 8.3). Atto565 NHS esters were resuspended in anhydrous DMSO (Sigma) and then aliquoted to and dried down into 0.25 µg/mL stocks under house vacuum and then stored at −20°C. Virus stock was added to dried Atto565 NHS ester and incubated at room temperature for 1 hour in the dark. After 1 hour, Tris pH 8.0 to a final concentration of 200 mM was added to quench the reaction. Free label was then separated from labeled virus using a 0.5 mL Piece Zeba spin desalting column with a 40 kDa molecular weight cut off. Labeled virus was stored at 4°C and used within 1 week of labeling.

### Preparation of glass coverslips

VSV-SARS-CoV-2-Atto565 infection were performed on 25 mm #1.5 coverslips bound with polydimethylsiloxane (PDMS) with 3 mm (infection) or 5 mm (uptake) wells as previously described (*31*). After desired wells were punched, PDMS was attached to the top surface of a coverslips previously cleaned by sonication, first in isopropanol for 20 minutes and then in 0.5 M KOH for 20 minutes, followed by extensive washing in MilliQ water, followed by drying for 30 minutes at 90°C. The glass was then firmly pressed to the PDMS sheet, after exposure of the surfaces to air plasma in a PDC-001 plasma cleaner (Harrick Plasma) operating at 750 mtorr and 30W for 2 min. The bonded glass/PDMS was heated at 90°C for 20 min, followed by sterilization by incubation with 70% ethanol for 10 min immediately before plating of cells.

For the single particle tracking LLSM experiments, round #1.5 glass coverslips of 6 mm in diameter) were cleaned by sequential 20 min sonication in toluene, dichloromethane, ethanol, and then water followed by storage in 70% ethanol until used for cell plating.

### Single molecule Atto565 dye calibration

VSV-SARS-CoV-2-Atto565 was plated for 10 minutes on glass coated with 0.1 mg/mL of poly-D-lysine for 30 minutes (then extensively washed in PBS) and unbound virus removed by 3 consecutive washes with PBS. Atto 565 signal per virion was determined by Spinning disk was imaged by taking 20 planes in z spaced 0.27 µm apart at 50 ms exposures. The number of Atto 565 dyes per virus was calibrated by increasing the exposure (while maintaining the same laser power) of the virus to 1 second and bleaching the Atto 565 dye. The bleach kinetics of Atto565 was fit using a step detecting code in MATLAB (*34*) and the intensity of the last step bleach steps of single particles was used to determine the intensity of a single Atto565 dye. The linear relationship between intensity and exposure allowed for this intensity to be used to determine the number of dyes attached to each virus.

### Infection assays for VSV-SARS-CoV-2

Infection assays were done with cells plated one day before the experiment and used for the infection assays with a final density of ~ 80% confluency as previously described (*31*). VSV-SARS-CoV-2-Atto 565 was incubated with the indicated concentration of antibody for 1 hour at 37°C. Virus was then added to the cells with indicated concentration of antibody for 1 hour, followed by 3 washes with cell culture media (DMEM, 10% FBS, 25 mM HEPES, pH 7.4), and then incubation for 7 hours at 37°C and 10% CO_2_ in DMEM containing 25 mM HEPES, pH 7.4.

At the end of the 7-hr incubation, cells were treated for 30 sec with 5 µg/mL WGA-Alexa647 in PBS to define the cell outline, then fixed with 4% PFA in PBS for 15 minutes at room temperature and imaged within 24 hours using a spinning-disk confocal microscope equipped with a 40x oil NA=1.4 objective (1 pixel = 0.33 µm) to acquire random fields, each containing a Z-stack of 20 consecutive optical planes taken 1 µm apart. Cells were scored as infected if the cytosolic eGFP fluorescence was 1.4 times that of the background of uninfected cells from the control experiment without virus.

### Lattice light sheet microscopy

Cells were plated onto 5 mm coverslips in a 35 mm culture dish at 60% confluence the day prior to each experiment. Approximately 10 µl of a solution containing 2.5 µg/mL viral RNA of VSV-SARS-CoV-2-Atto565 (MOI ~2) was added to coverslips placed on a piece of parafilm in a 10 cm petri dish for 4 minutes in DMEM with 25 mM HEPES at pH 7.4. Experiments with SP1-77 or VHH7-5-82 virus was pre-incubated with 500 ng/mL antibody at 37°C for 1 hour then 10 µL of VSV-SARS-CoV-2/antibody solution was added to the coverslip. Wetted Chem wipes were place around the parafilm in the Petri dish to help maintain humidity of the sample. The cells were imaged in phenol red free media, (FluoBright) supplemented with 5% FBS and 25 mM HEPES, pH 7.4. The samples were volumetrically imaged as a time series using a dithered multi-Bessel lattice light sheet using a sample scan with 0.5 µm spacing between each plane every 4 seconds. Cells were kept at 37°C and 5% CO2 for the duration of the experiment and soluble Alexa 549 fluorescent dye at 100 nM added to incubation bath to outline cells in experiments.

### Single virus tracking and image analysis

The 3D stacks from the LLSM experiments were deskewed and the diffraction limited spots were detected and tracked using an automated detection algorithm that uses least-squares numerical fitting with a model of the microscope PSF approximated by a 3D Gaussian function and implemented in MATLAB previously developed (*35*). Estimated fluorescence intensities associated with each spot were calculated from the corresponding amplitudes of the fitted 3D Gaussian and compared to single virus bound to poly-D-lysine coated glass imaged under the same condition as the experiments. Tracks of virus were then exported into a custom-made program written in LabView for visualizing trajectories (*32*). Each virus trajectory was visually examined for localization to the cell surface, internalization into the cell, and spreading of the Atto565 dye fluorescence. The virus is 80-100 nm, which makes it a diffraction limited object. The spot limited by the wavelength of the dye being emitted is 300 nm. The endosome is 600 nm and larger. The fusion of the viral envelope to the endosomal membrane is observed by a spreading of the fluorescent signal from a point spread limited object to 500-1000 nm over the endosomal membrane.

### S1 dissociation reconstitution

The amount of Atto565 signal per virus was determined using single molecule calibration described above under different treatments to determine the proteolytic and ACE2 requirement for S1 dissociation. VSV-eGFP-SARS-CoV-2-Atto565 was incubated with 1 µg/mL trypsin for 30 minutes at 37°C in PBS then the reaction was stopped by the addition of 10 µM Aprotinin final concentration. Virus infection was previously confirmed to no longer require host cell proteases after trypsin cleavage (*32*). Virus was then bound to poly-D-lysine coated glass single molecule calibration determine no different in fluorescence between trypsin and non-trypsin treated virus. Incubating virus with ACE2 for 10 min. at 37°C after trypsin cleavage treatment caused a ~75% decrease in the number of Atto565 molecules per virus. While treatment with 100 ng/mL of SP1-77 IgG or SP1-77 Fab after trypsin cleavage prior to ACE2 binding prevented this loss of signal.

---

**Figure S1.**
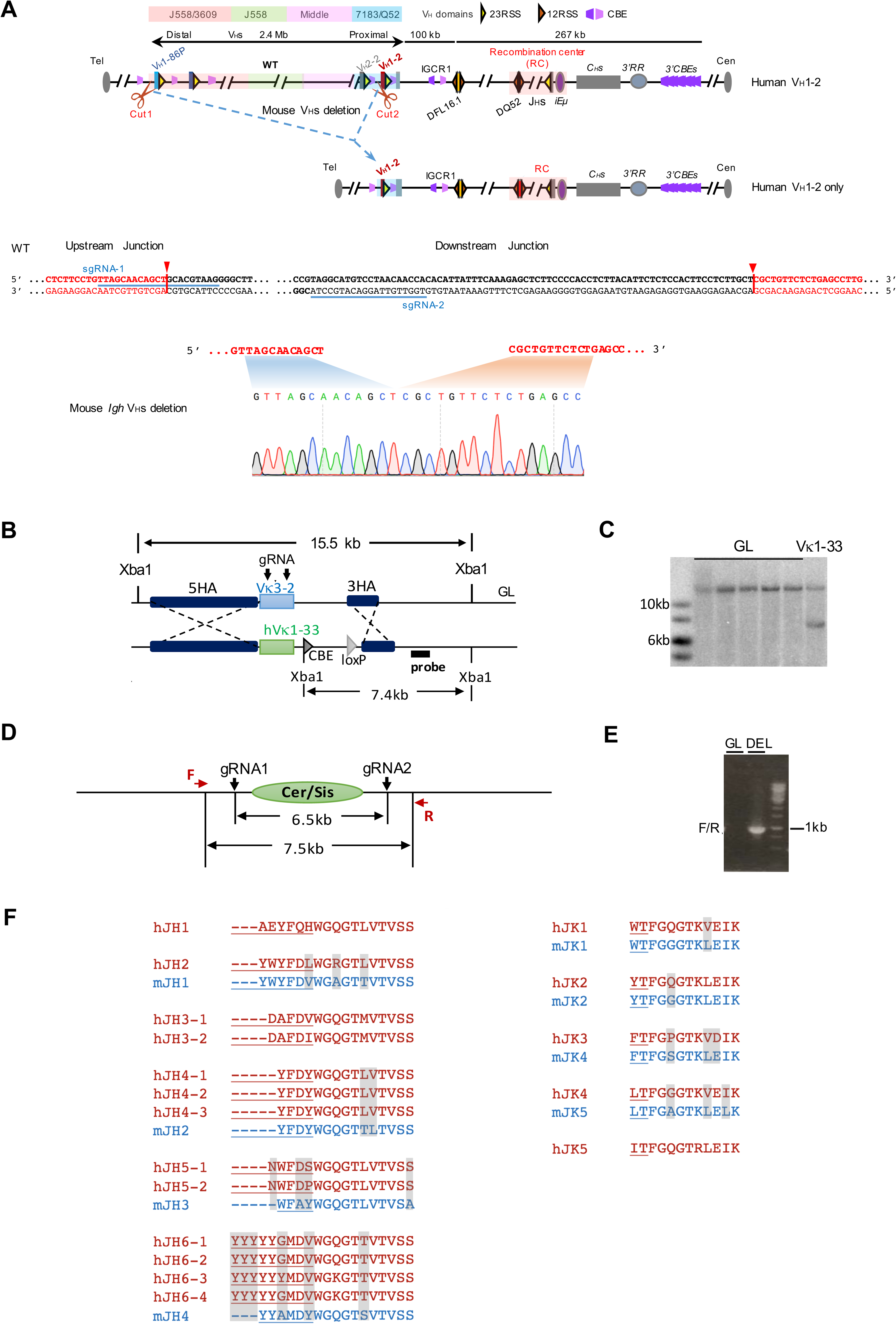
Generation of the V_H_1-2/Vκ1-33-rearranging mouse model. (**A**) The diagram illustrates the deletion of 2MB upstream region of human V_H_1-2 that contain all mouse V_H_s. The deletion was mediated by two gRNAs (green). The deleted allele was screened by PCR. The sequencing result was shown in the bottom. (**B**) The diagram, not drawn to scale, illustrates the restriction digests and Southern probe that were used to differentiate the region before (GL) and after Vκ1-33 replacement. (**C**) Southern analysis of positive Vκ1-33 ES clones that showed in (**B**). (**D**) The diagram illustrates the deleted region of Cer/sis regulatory element targeted by two gRNAs and PCR primers that were used to identify the deleted clones. (**E**) PCR analysis of ES clones with Cer/sis deletion. (**F**) Similarities of amino acid sequences between mouse Js and human Js. Human Js are shown in red, and mouse Js are shown in blue. The alignments between human Js and mouse Js were mainly based on the sequences underlined at 5’ end that contribute to the CDR3 sequence. The different amino acids between mouse Js and human Js are filled in grey.

**Figure S2.**
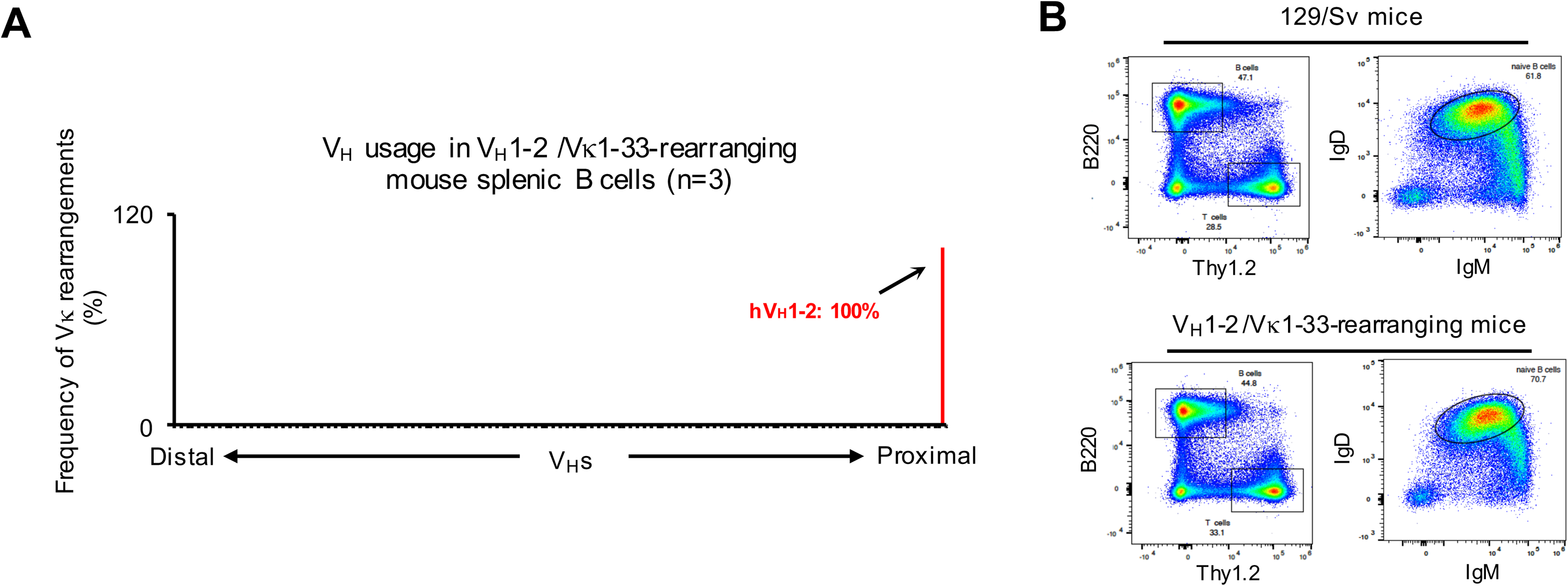
Characterization of the V_H_1-2/Vκ1-33-rearranging mouse. (**A**) HTGTS-rep-seq analysis of V_H_ usage in V_H_1-2^mVHΔ/IGCR1Δ^-rearranging mouse splenic B cells. The x axis lists all functional V_H_s from the distal to the *D*-proximal end. The histogram displays the percent usage of each V_H_ among all productive V_H_DJ_H_ rearrangements. (**B**) FACS analyses of splenic B cell and T cell populations from wild-type 129/Sv and V_H_1-2/Vκ1-33-rearranging mice. They are representative of three biological replicates.

**Figure S3.**
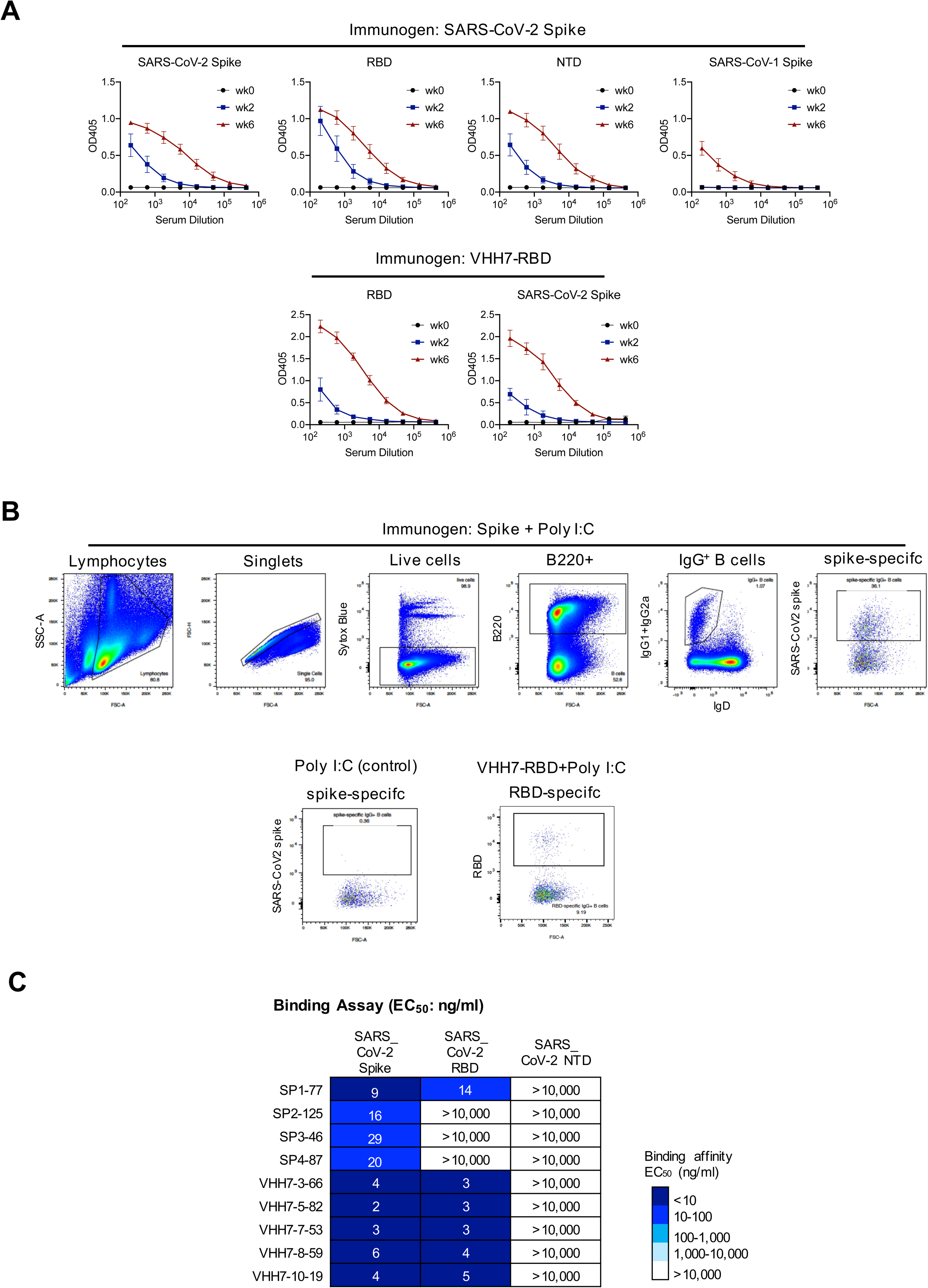
Immunizing the mouse model with SARS-CoV-2 spike or RBD elicited V_H_1-2/Vκ1-33 antibodies. (**A**) Binding curves showing reactivity of sera to SARS-CoV-2 related antigens. The upper panel shows the sera from the SARS-CoV-2 Spike immunized mice at week 0, 2 and 6. The bottom panel shows the sera from VHH7-RBD immunized mice. Data were mean ± SD of three mice. (**B**) Gating strategy for single cell sorting of SARS-CoV2 spike-specific or RBD-specific IgG^+^ B cells after immunization. The upper panel showed the mice immunized with SARS-CoV2 spike plus poly I:C adjuvant. The bottom panels showed the mice immunized with poly I:C (control) or VHH7-RBD plus poly I:C adjuvant. (**C**) Table shows the binding affinities of nine mAbs to the SARS-CoV-2 spike protein, RBD, NTD. Data are representative of two independent experiments. EC_50_ values were color-coded based on the key shown at the right.

**Figure S4.**
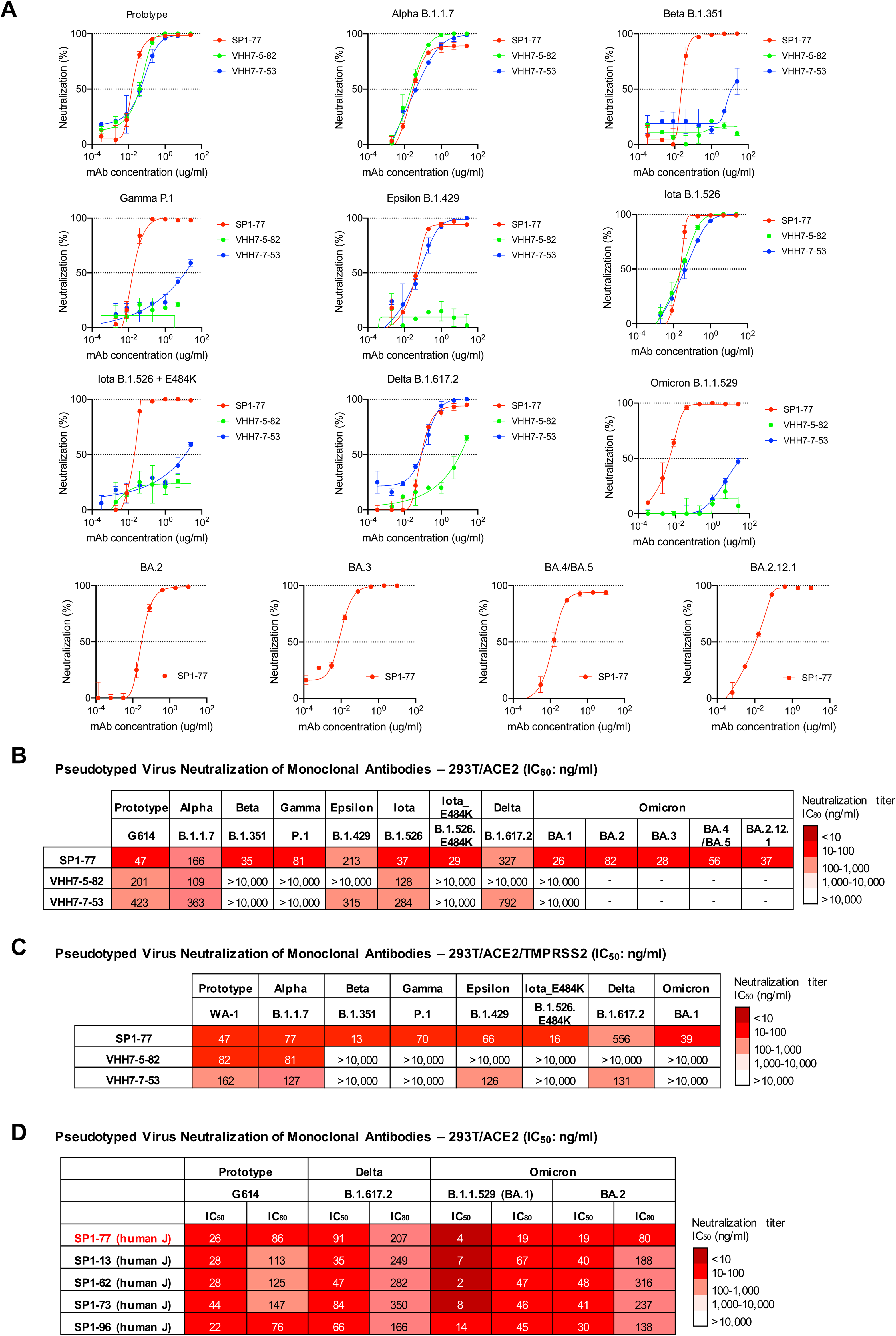
Neutralization data of three mAbs against SARS-CoV-2 pseudoviruses. (**A**) Neutralization Curves of SARS-CoV-2 pseudoviruses displaying S proteins by three V_H_1-2/Vκ1-33 mAbs. Data were done in 293T/ACE2 cells by Montefiori et al. and also shown in Fig. 3C and fig. S4B. Data are representative of one independent experiments with two technical replicates. (**B**) Table shows the neutralization activity - IC_80_ of three mAbs against VOCs and VOIs in pseudovirus neutralization assays. Experiments were done in 293T/ACE2 cells and also shown in Fig. 3C and fig. S4A. Data are representative of 2 biologically independent experiments for most VOCs and VOIs, but one experiment for BA.2 and BA.3. Each independent experiment contains 2 technical replicates. IC_80_ values are color-coded based on the key shown at the right. (**C**) Table shows the neutralization activities of three mAbs against VOCs and VOIs in pseudovirus neutralization assays. These assays were done independently in 293T/ACE2/TMPRSS2 cells. Compared to the neutralization data done in 293T/ACE2 cells, IC_50_ values are at the similar levels for all VOC and VOI except Delta which was neutralized less potently. Data are representative of one independent experiment with two technical replicates. IC_50_ values are color-coded based on the key shown at the right. (**D**) Table shows the neutralization activities of humanized SP1-77 and the other 4 antibodies in SP1 clonal lineage against VOCs and VOIs in pseudovirus neutralization assays. These assays were done in 293T/ACE2 cells. Data are representative of two independent experiments. Each independent experiment contains 2 technical replicates. IC_50_ values are color-coded based on the key shown at the right.

**Figure S5.**
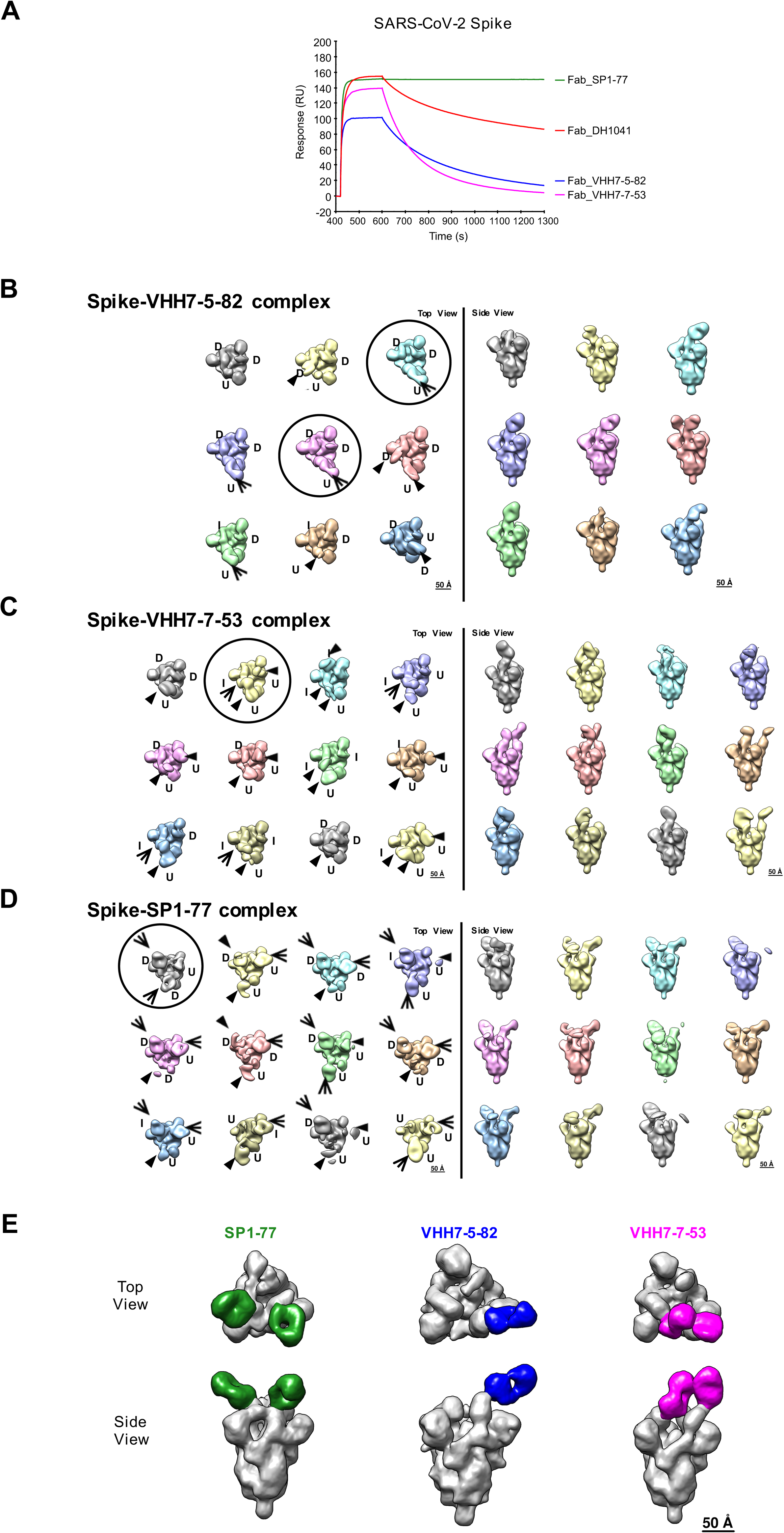
Negative stain electron microscopy analysis of S complexes with three V_H_1-2/Vκ1-33 neutralizing antibodies. (**A**) SPR data measuring the binding affinity between monoclonal Fabs and SARS-CoV-2 spike proteins. Each coronavirus spike (S) protein was captured on Streptavidin sensor chip and Fabs were screened at 50nM. A blank streptavidin surface along with blank buffer binding were used for double reference subtraction to account for non-specific protein binding and signal drift. Data is representative of two independent experiments. (**B**) A 3D classification of the spike-VHH7-5-82 complex data, with 158,101 particles sorted into 9 classes, shown in top and side views. (**C**) A 3D classification of the spike-VHH7-7-53 complex data, with 187,262 particles sorted into 12 classes, shown in top and side views. (**D**) A 3D classification of the spike-SP1-77 complex data, with 230,136 particles sorted into 12 classes, shown in top and side views. In the top view of **B-D**, arrows indicate Fabs, and arrowheads indicate weak or partial Fab density; RBDs were visually assessed and assigned as up (U), down (D) or intermediate (I); and the circles indicate the selected (or combined) classes used for the final 3D reconstructions shown in **E**. (**E**) Final 3D reconstructions of Fab-S complexes shown in top view and side view with the S in gray and the Fabs colored (SP1-77: green; VHH7-5-82: blue; VHH7-7-53: magenta).

**Figure S6.**
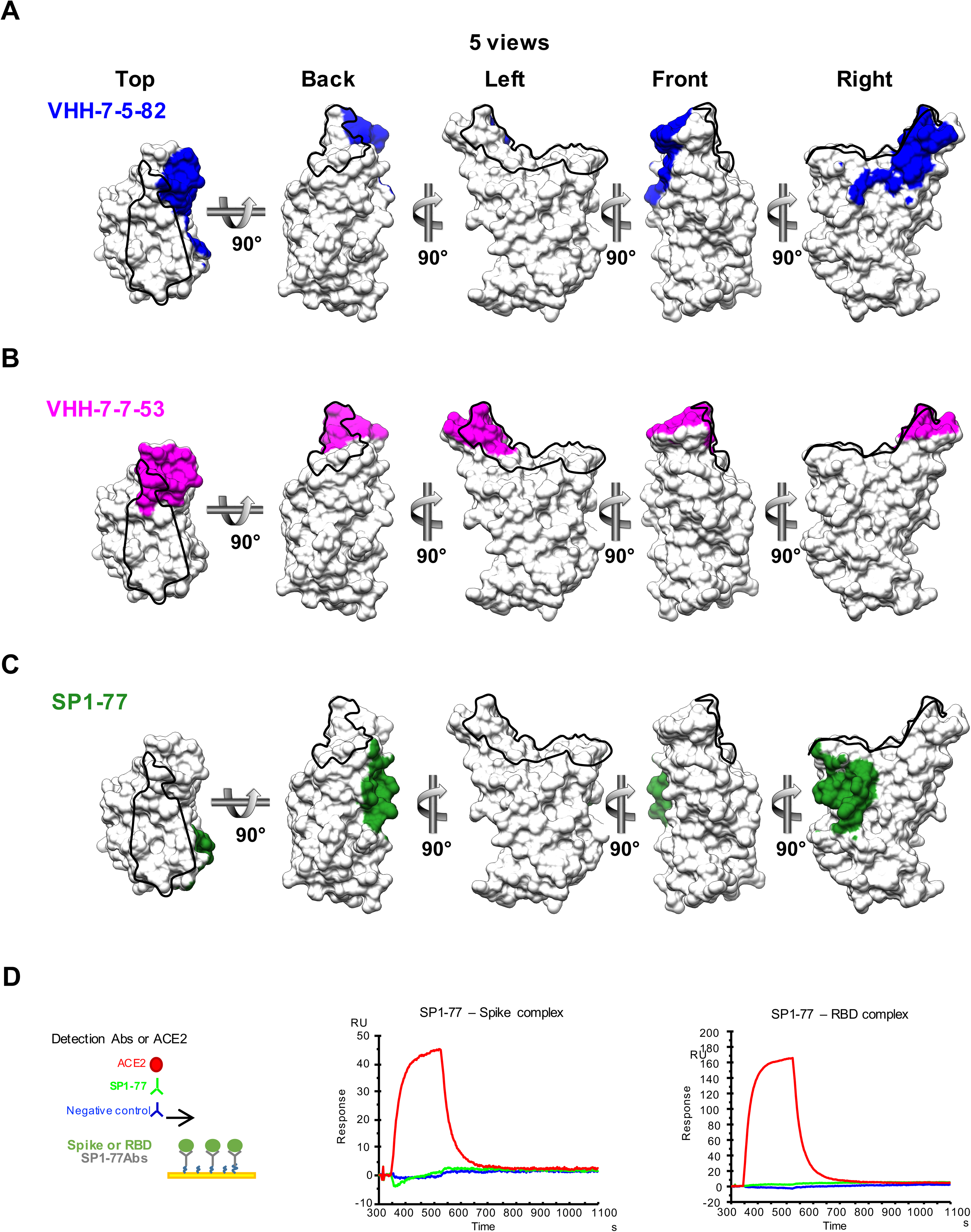
Three V_H_1-2/Vκ1-33 neutralizing antibodies bind to distinct regions of RBD. (**A-C**) Binding footprint of Fabs of three V_H_1-2/Vκ1-33 neutralizing antibodies, VHH7-5-82 (**A**), VHH7-7-53 (**B**), and SP1-77 (**C**) shading on the RBD in surface representation. The RBD surface was represented in 5 views (top, back, left, front and right). Black outline indicates the ACE2 RBM. RBD was shown as gray. (**D**) SP1-77 can not block ACE2 binding to the spike protein and RBD. The left panel showed the diagram of SPR assay. The middle panel showed the binding of ACE2, SP1-77 antibody and negative control (red, green and blue, respectively) to preformed SP1-77 - spike complex. The right panel showed the binding of ACE2, SP1-77 antibody and negative control (red, green and blue, respectively) to preformed SP1-77 - RBD complex.

**Figure S7.**
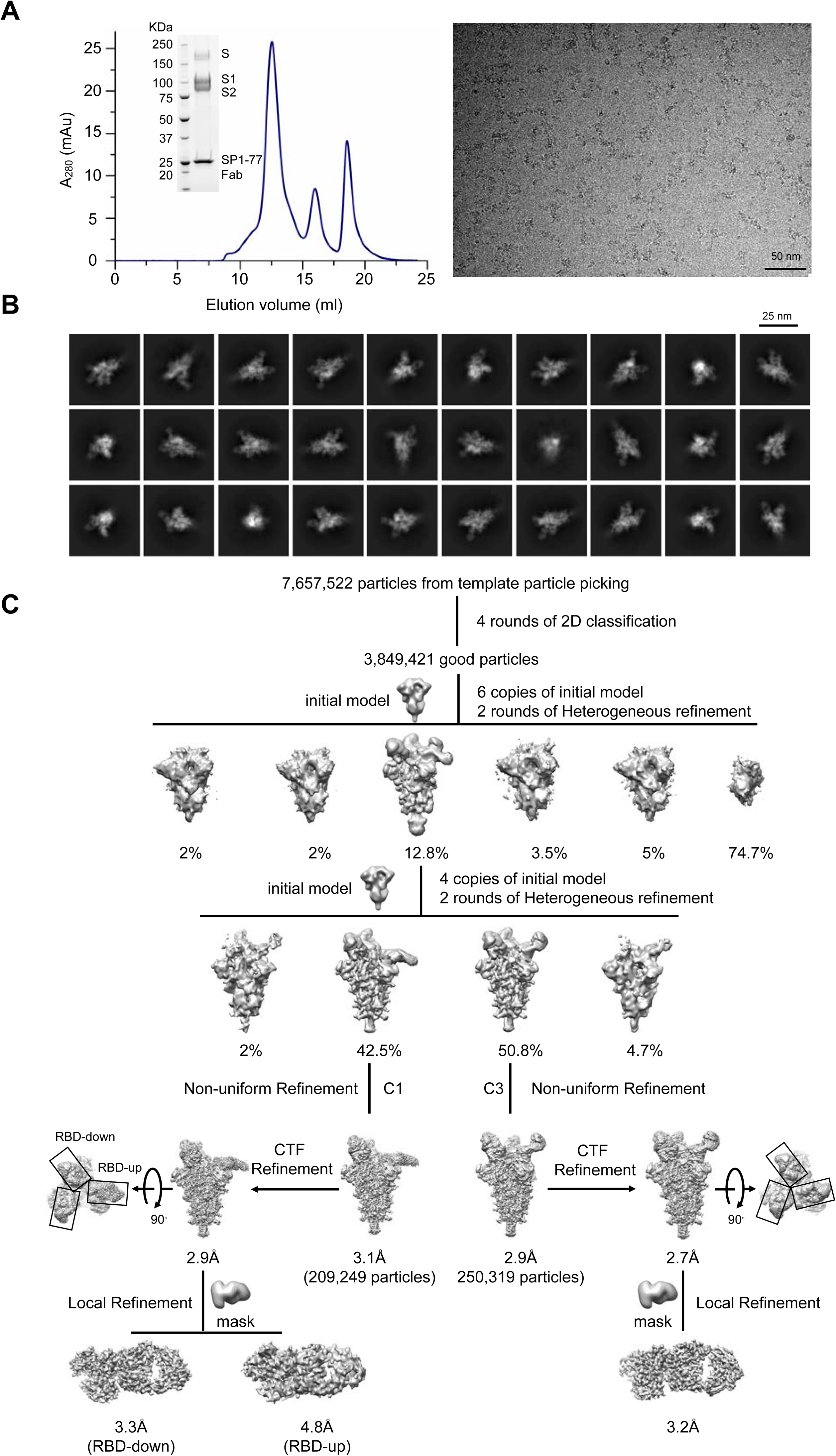
Purification of the full-length G614 S trimer/SP1-77 Fab complex and cryo-EM analysis. (**A**) Left, the complex of the full-length G614 S trimer and SP1-77 Fab was resolved by gel-filtration chromatography on a Superose 6 column. First peak, the S trimer-Fab complex used for the cryo-EM study; second peak, the dissociated postfusion S2 trimer mixed with the monomeric S1 in complex with Fab; and third peak, free Fab. Inset, peak fractions were analyzed by Coomassie stained SDS-PAGE. Right, representative motion-corrected micrograph of the vitrified complex purified by gel-filtration chromatography. (**B**) 2D class averages (box dimension: 396Å) of the cryo-EM images of the G614 S trimer/SP1-77 Fab complex from cryoSPARC. (**C**) Data processing workflow for structure determination.

**Figure S8.**
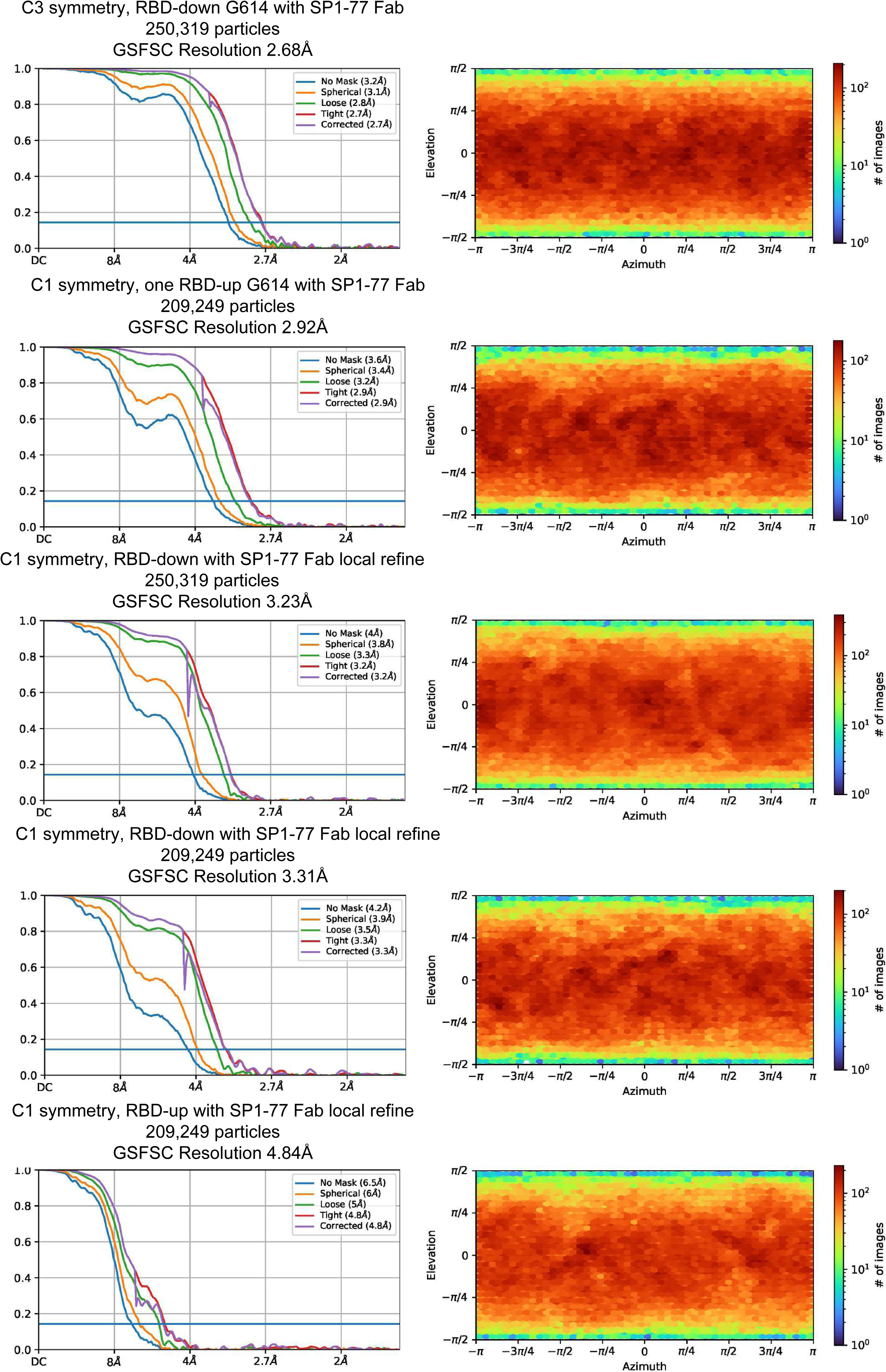
Cryo-EM structure validation of the G614 S trimer/SP1-77 Fab complex. FSC curves and the viewing direction distribution plot for the S trimer-Fab complex. From top to bottom: the complex of the G614 S trimer in the three-RBD-down conformation and three SP1-77 Fabs; the complex of the G614 S trimer in the one-RBD-up conformation and three SP1-77 Fabs; masked local refinement of the Fab in complex with the RBD from the G614 S trimer in the three-RBD-down conformation; masked local refinement of the Fab in complex with the RBD in the up conformation from the G614 S trimer in the one-RBD-up state; masked local refinement of the Fab in complex with the RBD in the down conformation from the G614 S trimer in the one-RBD-up state.

**Figure S9.**
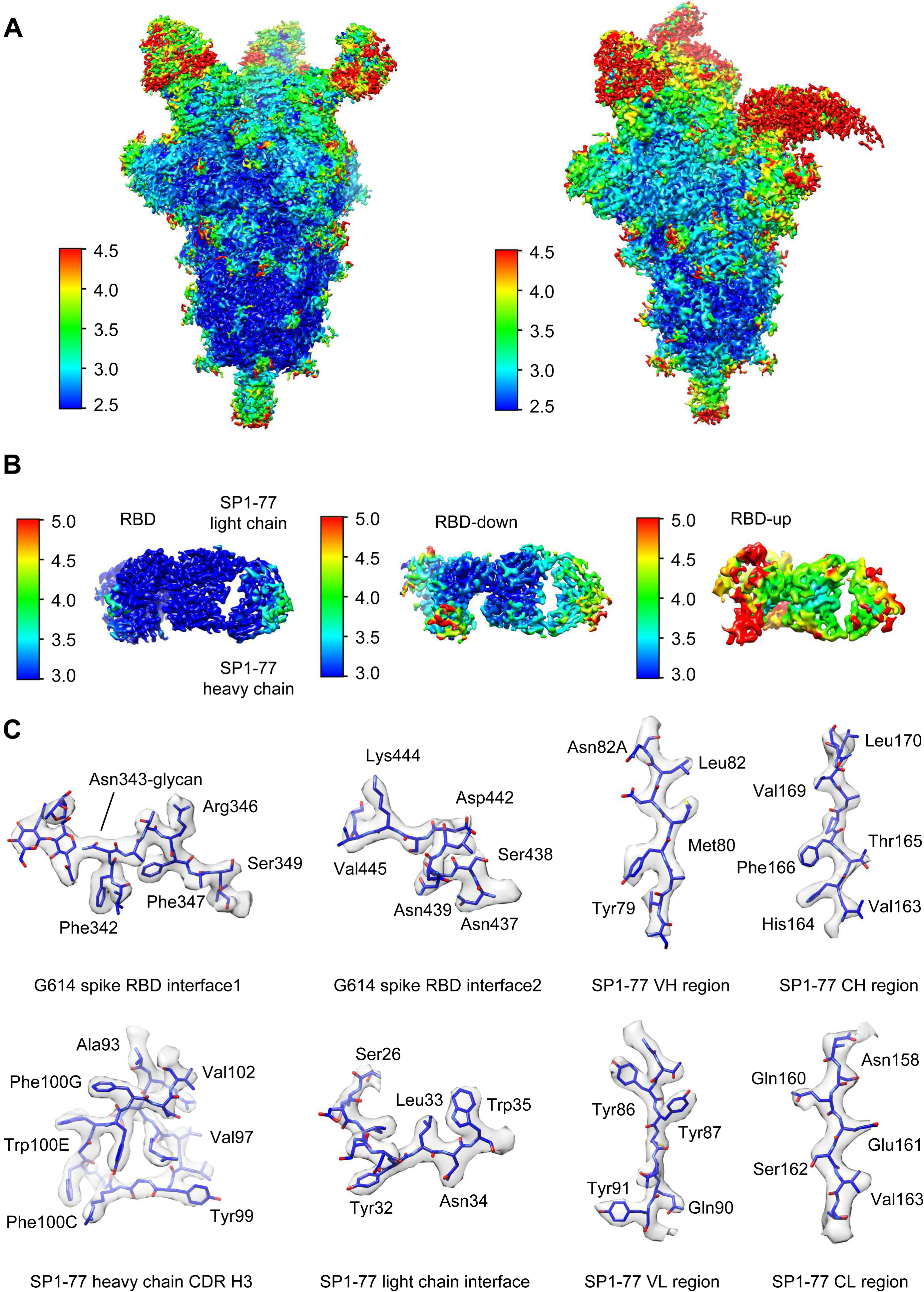
Analysis of the the G614 S trimer/SP1-77 Fab complex structure. (**A**) 3D reconstructions of SP1-77 Fab in complex with the G614 S trimer in the three-RBD-down and one-RBD-up conformations, respectively, are colored according to local resolution estimated by cryoSPRAC. (**B**) 3D reconstructions from masked local refinement of the Fab in complex with a single RBD from the G614 S trimer in the three-RBD-down conformation (left), or the RBD in the up conformation from the G614 S trimer in the one-RBD-up state (middle) or the RBD in the down conformation from the G614 S trimer in the one-RBD-up state (right) are colored according to local resolution estimated by cryoSPARC. (**C**) Representative density in gray surface from EM maps with a resolution better than 3.3Å, primarily focusing on the SP1-77 Fab and its binding interface with the RBD of the G614 S trimer.

**Figure S10.**
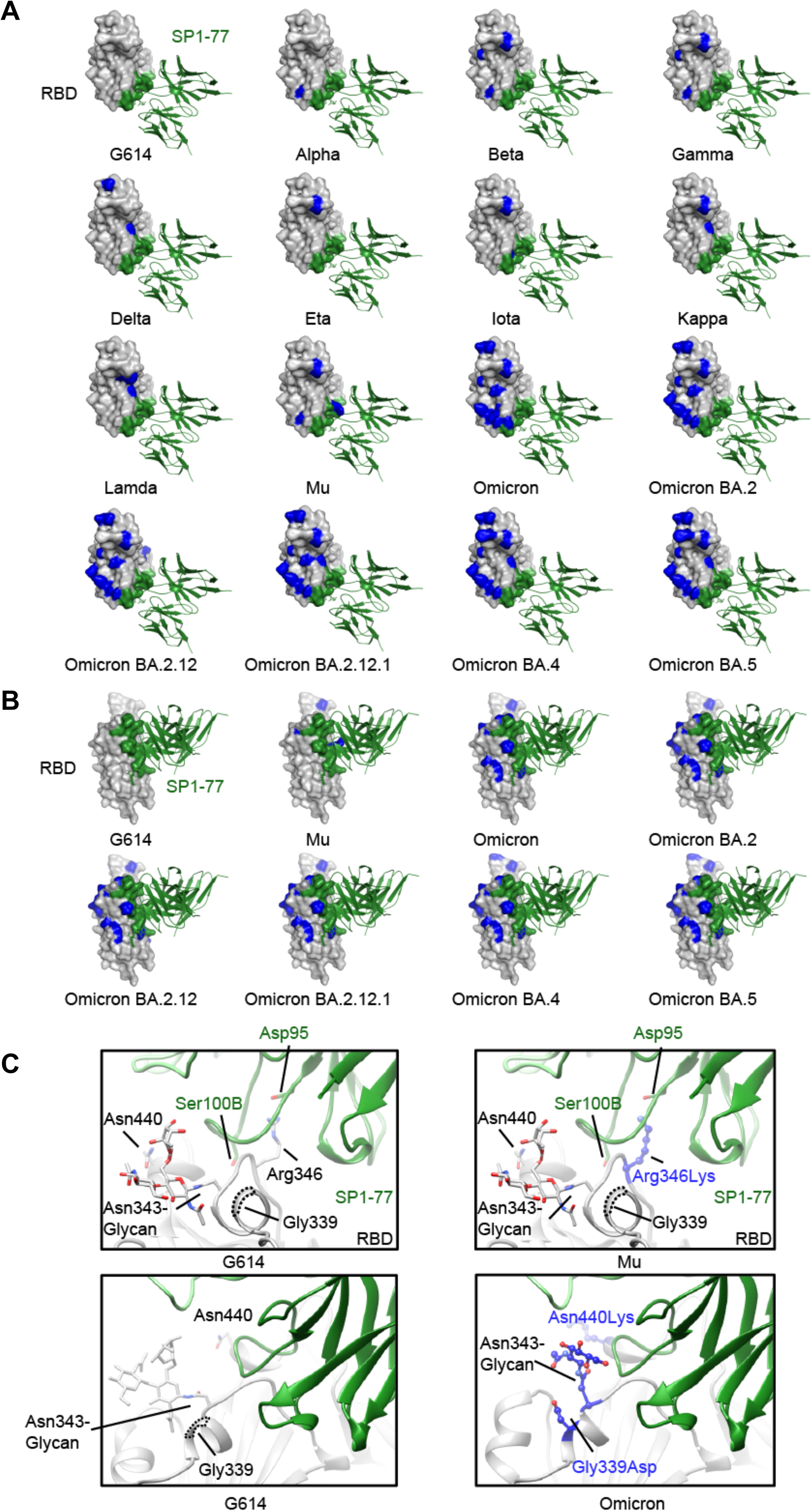
Modeled SP1-77 binding site on various SARS-CoV-2 variants. (**A**) The potential footprint of SP1-77 on the modeled RBDs from different SARS-CoV-2 variants in a top view. The RBD is shown in surface representation in gray with the SP1-77 footprint highlighted in green and the mutations in each variant in blue. The Fv region of SP1-77 is shown in ribbon diagram in green. Most mutations on spike variants are not located at the SP1-77 footprint. (**B**) Side view of a selected panel from **A**. (**C**) Structural comparison of the SP1-77 binding interface among the RBDs of wildtype G614, Mu and Omicron variants. The conservative nutation R343K in Mu preserves the salt bridge between the residue 343 in the RBD and the SP1-77 D99. The mutations in Omicron variant reconfigures the local conformation near the N343-glycan, which is on the edge of SP1-77 footprint. The RBD is colored in gray, the SP1-77 heavy chain in green and the light chain in light green. Mutations in the Mu and Omicron variants are shown in stick and ball model in blue.

**Figure S11.**
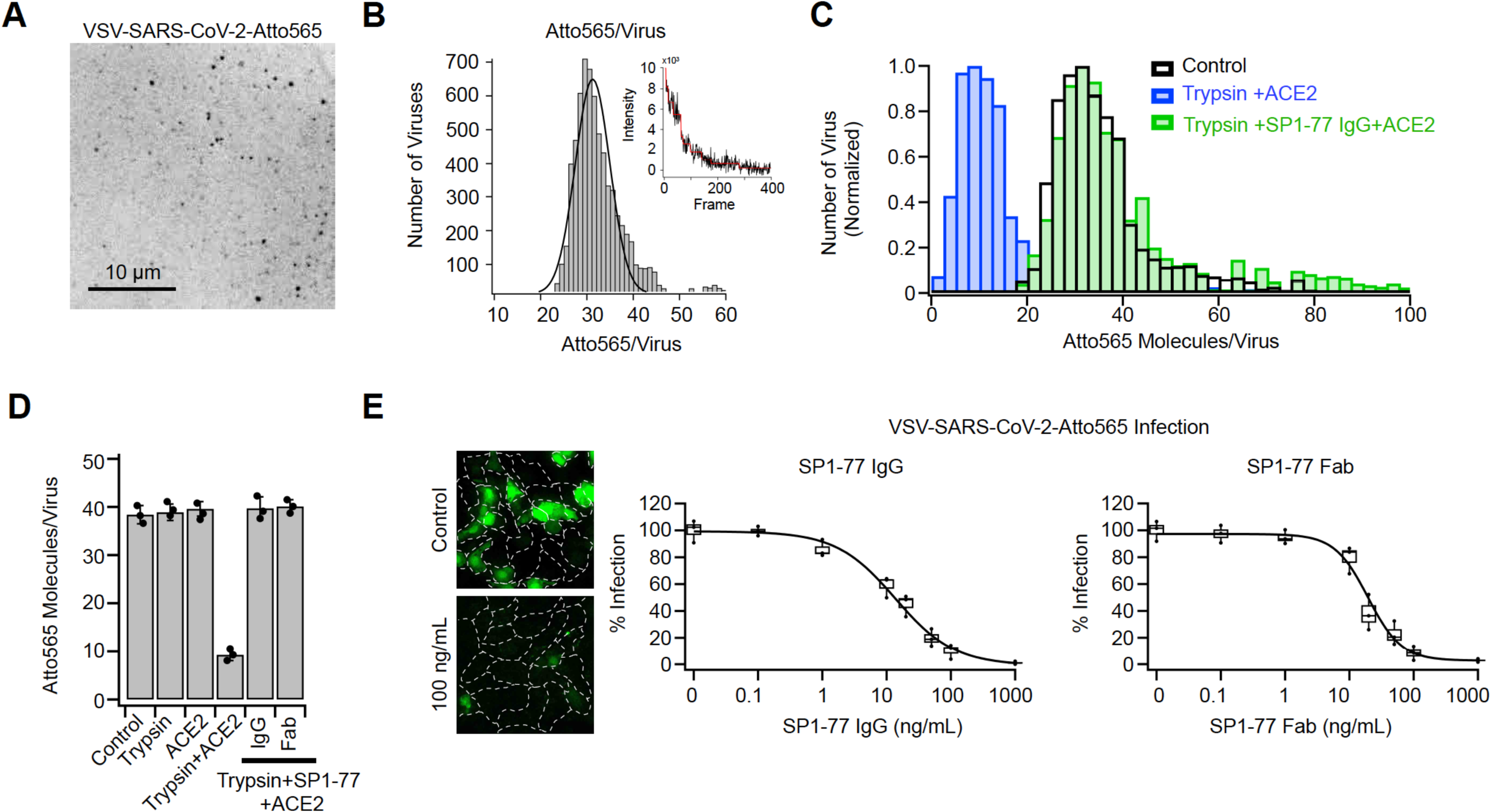
Single molecule calibration of VSV-SARS-CoV-2-Atto565 virus. (**A**) Example image of Atto 565 fluorescence of virus adsorbed for 10 minutes on a glass coverslip coated with poly-D-lysine coated slides and imaged and imaged with spinning disc confocal microscopy. Photobleaching (B), used to determine the intensity of single Atto 565 dye intensities allowed to establish that ~20-40 dyes were attached to each virus. (**C**) Histogram and (**D**) data summary of the number of Atto 565 molecules on VSV-SARS-CoV-2-Atto 565 determined by single molecule counting. Treatments prior to adsorption included none (control), incubation with 1 µg/mL trypsin for 30 min at 37°C, treatment without or with trypsin followed by an incubation with 0.5 µM of recombinant ACE2 for 10 min at 37°C, or treatment with trypsin then incubated with 100 ng/mL SP1-77 IgG or SP1-77 Fab for 1 hour at 37°C followed by incubation with ACE2. (**D**) Statistics are averages of peak distribution determined by a Gaussian fit of 3 independent experiments. (**E**) Infection assays showing neutralization of VSV-SARS-CoV-2-Atto 565 infection with SP1-77 IgG or SP1-77 Fab. A soluble eGFP reporter genetically encoded into this VSV chimera allowed for infection to be determined by fluorescence imaging on a spinning disc confocal microscope as shown in the representative examples obtained 7 hr post infection. Cell outlines were obtained with a WGA-Alexa 647 membrane stain applied to cells immediately prior to fixation. Each condition was measured with 3 independent experiments.

**Figure S12.**
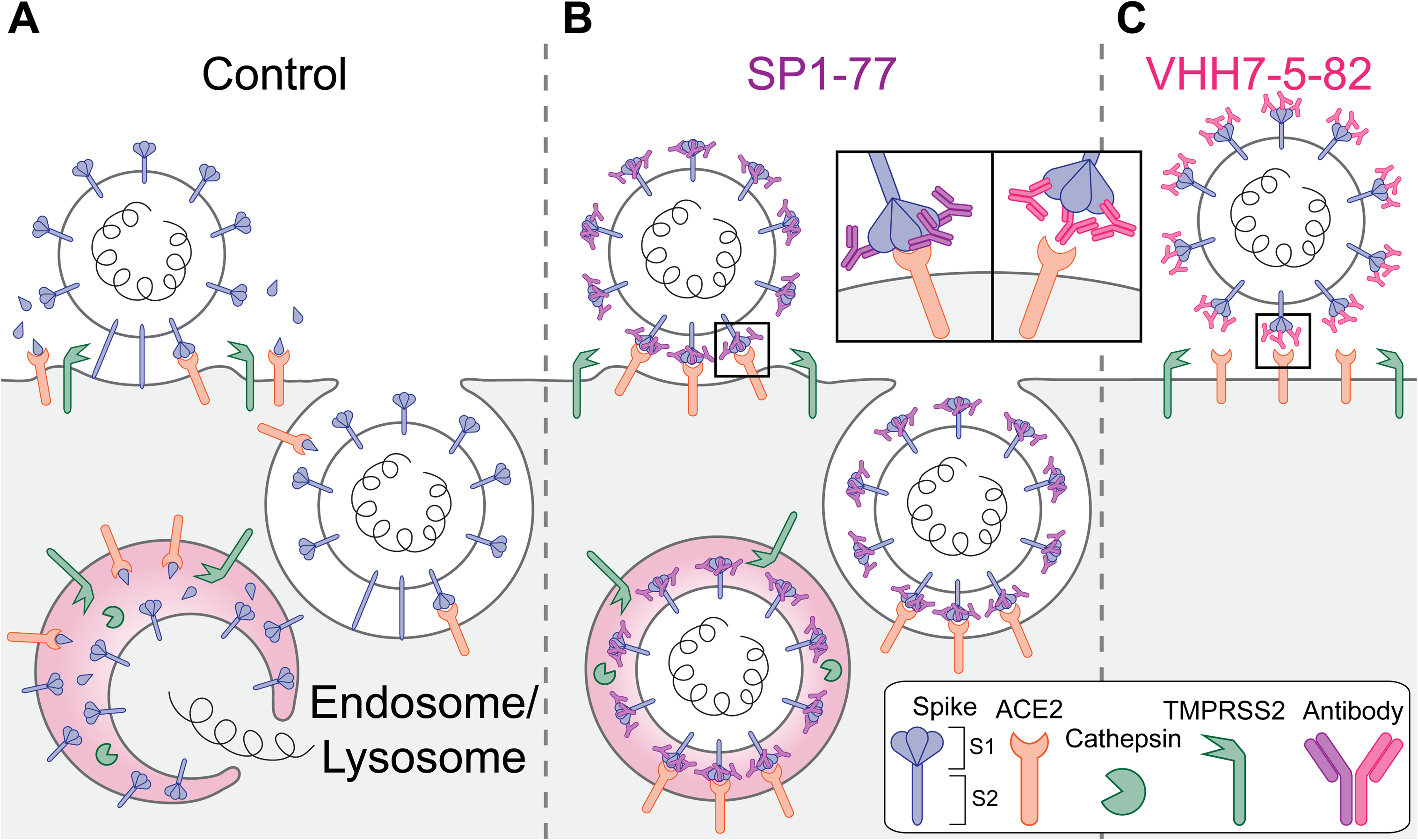
Schematic representation of mechanism of SP1-77 inhibition of SARS-CoV-2 infection. (**A**) Without antibody treatments, the spike protein on the viral surface binds to the ACE2 receptor on the infected cell surface. Membrane fusion is activated either by TMPRSS2 protease on the cell membrane or by cathepsin L protease following endocytosis. Cleavage at the S2’ site by these proteases leads to dissociation of the S1 subunit, which exposes the fusion peptide on the S2, facilitating viral-host membrane fusion and viral entry into the infected cells. (**B**) Pre-treatment of the virus with SP1-77, a non-ACE2-blocking antibody, does not appreciably impact binding of viruses to the cell surface and their endocytosis. However, SP1-77 greatly inhibits the dissociation of S1 subunit, thereby, blocking activation of the fusion peptide and membrane fusion. (**C**) Pre-treatment of the virus with VHH7-5-82, an ACE2-blocking antibody, prevents binding of the virus to the cell surface.

**Figure S13.**
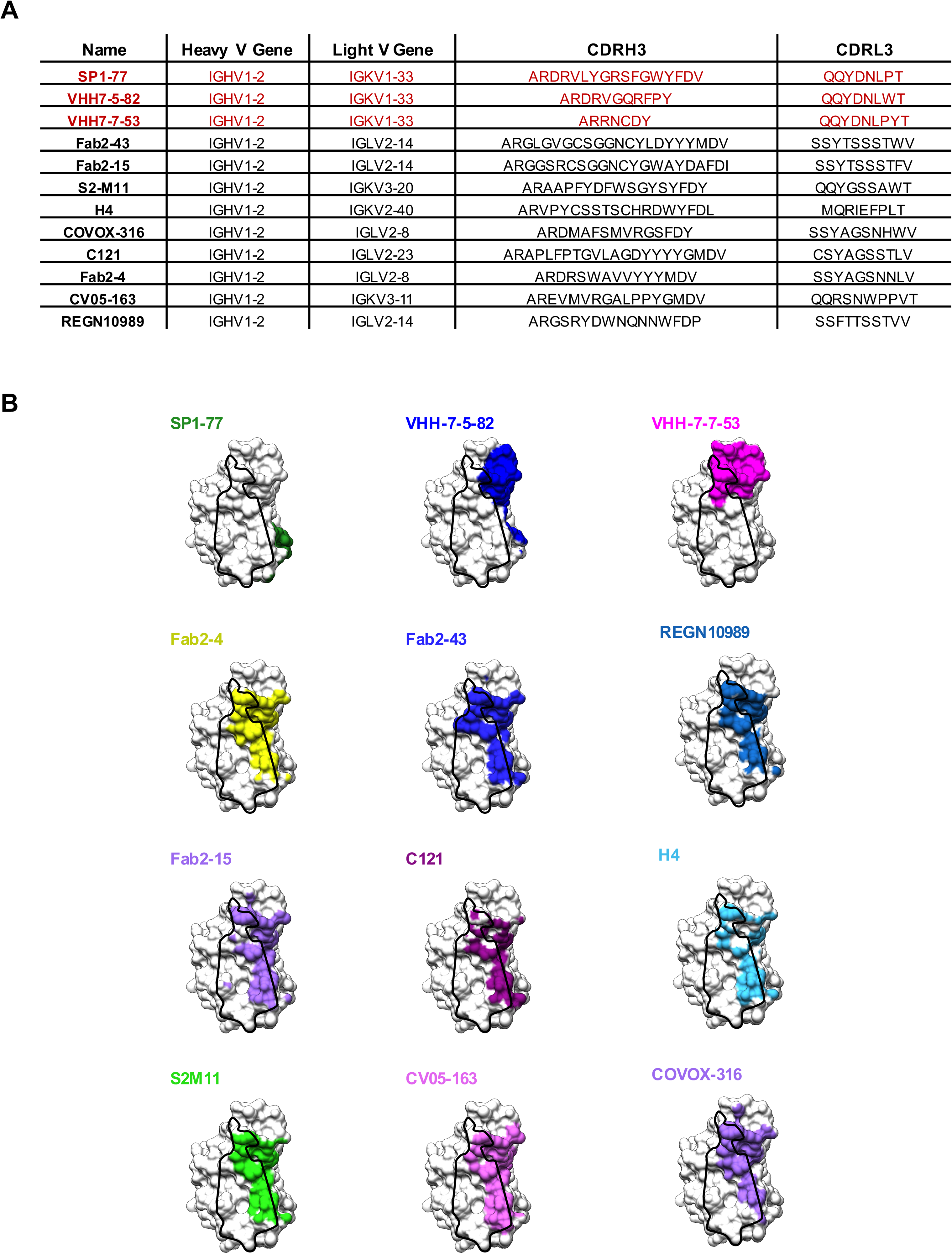
Epitope analysis and comparison of the previously characterized V_H_1-2-based mAbs. (**A**) Table shows a list of V_H_1-2-based mAbs for which high-resolution structures have been published with various HC CDR3s and LCs. The three antibodies identified in our mouse model were labeled in red. (**B**) Comparisons of footprints on RBD between our three antibodies and nine previously characterized V_H_1-2-based mAbs. Those nine previously characterized V_H_1-2-based mAbs show a similar RBD binding epitope that is distinct from where our three antibodies bind.

**Table S1:**
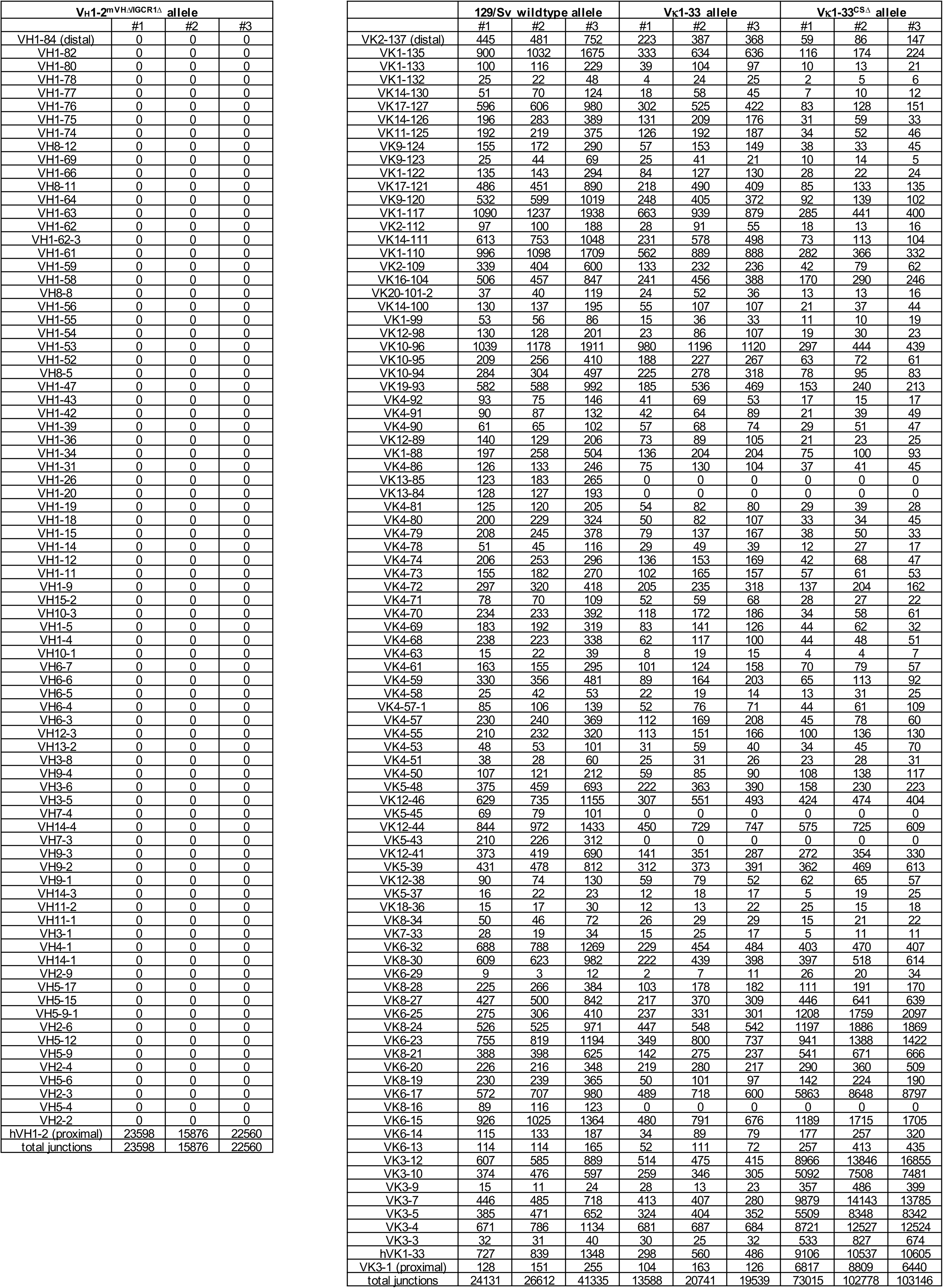
V usages in splenic B cells measured by HTGTS-Repertoire-seq.

**Table S2:**
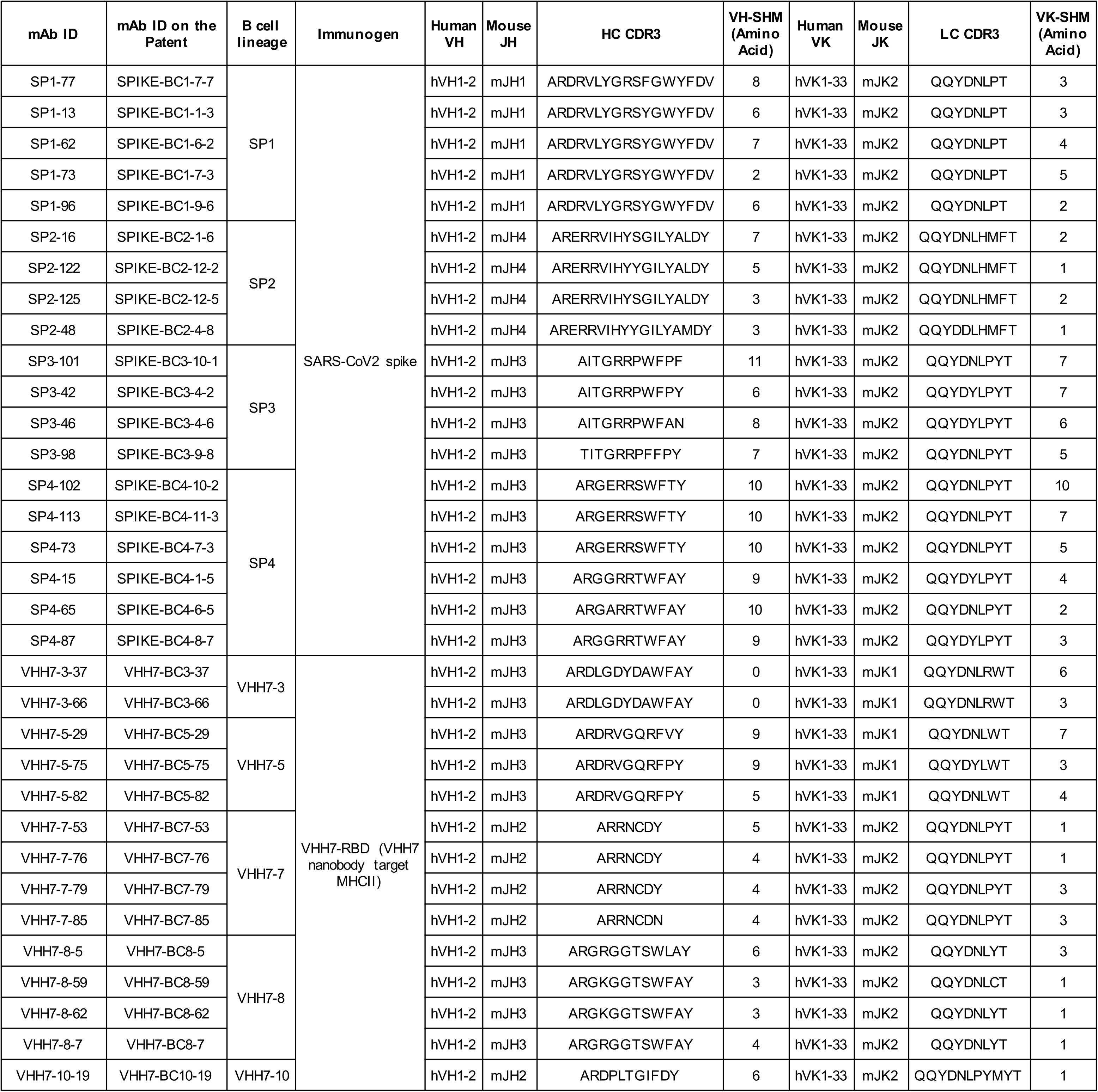
Sequence features of Anti-SARS-CoV-2 antibodies isolated from mouse model with single human V rearrangement.

**Table S3:** Cryo-EM statistics.

|  | Full-length SARS-CoV-2 G614 S:P1-37 Fab |  | RBD:P1-37 Fab |
| --- | --- | --- | --- |
| **Data collection and processing** |  |  |  |
| Microscope | Titan Krios |  | Titan Krios |
| Voltage(kV) | 300 |  | 300 |
| Detector | Gatan K3 |  | Gatan K3 |
| Magnification(nominal) | 105,000 |  | 105,000 |
| Energy filter slit width (eV) | 20 |  | 20 |
| Calibrated pixel size (Å/pix) | 0.83 |  | 0.83 |
| Exposure rate (e<sup>-</sup>/pix/sec) | 13.761 |  | 13.761 |
| Frames per exposure | 50 |  | 50 |
| Total electron exposure (e<sup>-</sup>/Å<sup>2</sup>) | 53.853 |  | 53.853 |
| Exposure per frame (e<sup>-</sup>/Å<sup>2</sup>) | 1.077 |  | 1.077 |
| Defocus range (µm) | -0.5,-2.2 |  | -0.5,-2.2 |
| Automation software | SerialEM |  | SerialEM |
| # of Micrographs used | 34,031 |  | 34,031 |
| Particles extracted | 7,657,522 |  | 7,657,522 |
| Particles after 2D classification | 3,849,421 |  | 3,849,421 |
| Class | RBD-down | RBD-up | RBD-down |
| Total # of refined particles | 250,319 | 209,249 | 250,319 |
| Symmetry imposed | C3 | C1 | C1 |
| Map resolution (Å) | 2.7 | 2.9 | 3.2 |
| FSC threshold | 0.143 | 0.143 | 0.143 |
| Map sharpening *B* factor (Å<sup>2</sup>) | -88.3 | -75.8 | -101.5 |
| **Refinement** |  |  |  |
| Initial model used (PDB ID) | 7KRQ, 6WS6,AlphaFold | 7KRR, 6WS6,AlphaFold | 6WS6, AlphaFold |
| Model composition |  |  |  |
| Non-hydrogen atoms |  |  |  |
| Amino acids | 4674 | 4619 | 635 |
| Glycans | 54 | 57 | 1 |
| RMSD bonds (Å) | 0.014 | 0.013 | 0.012 |
| RMSD angles (°) | 1.86 | 1.84 | 1.87 |
| Mean B-factors |  |  |  |
| Amino acids | 41 | 66 | 71 |
| Glycans | 56 | 94 | 86 |
| Ramachandran |  |  |  |
| Favored (%) | 93.04 | 93.36 | 93.16 |
| Allowed(%) | 5.86 | 5.70 | 5.41 |
| Outliers(%) | 1.10 | 0.94 | 1.43 |
| Rotamer outliers (%) | 2.59 | 2.22 | 2.20 |
| Clash score | 4.66 | 5.04 | 5.28 |
| C-beta outliers (%) | 0.53 | 0.39 | 0.68 |
| CaBLAM outliers (%) | 2.63 | 2.43 | 2.41 |
| CC (mask) | 0.79 | 0.77 | 0.87 |
| CC (volume) | 0.74 | 0.74 | 0.85 |
| MolProbity score | 2.00 | 1.96 | 1.98 |
| EMRinger score | 4.29 | 3.76 | 3.81 |

**Table S4:**
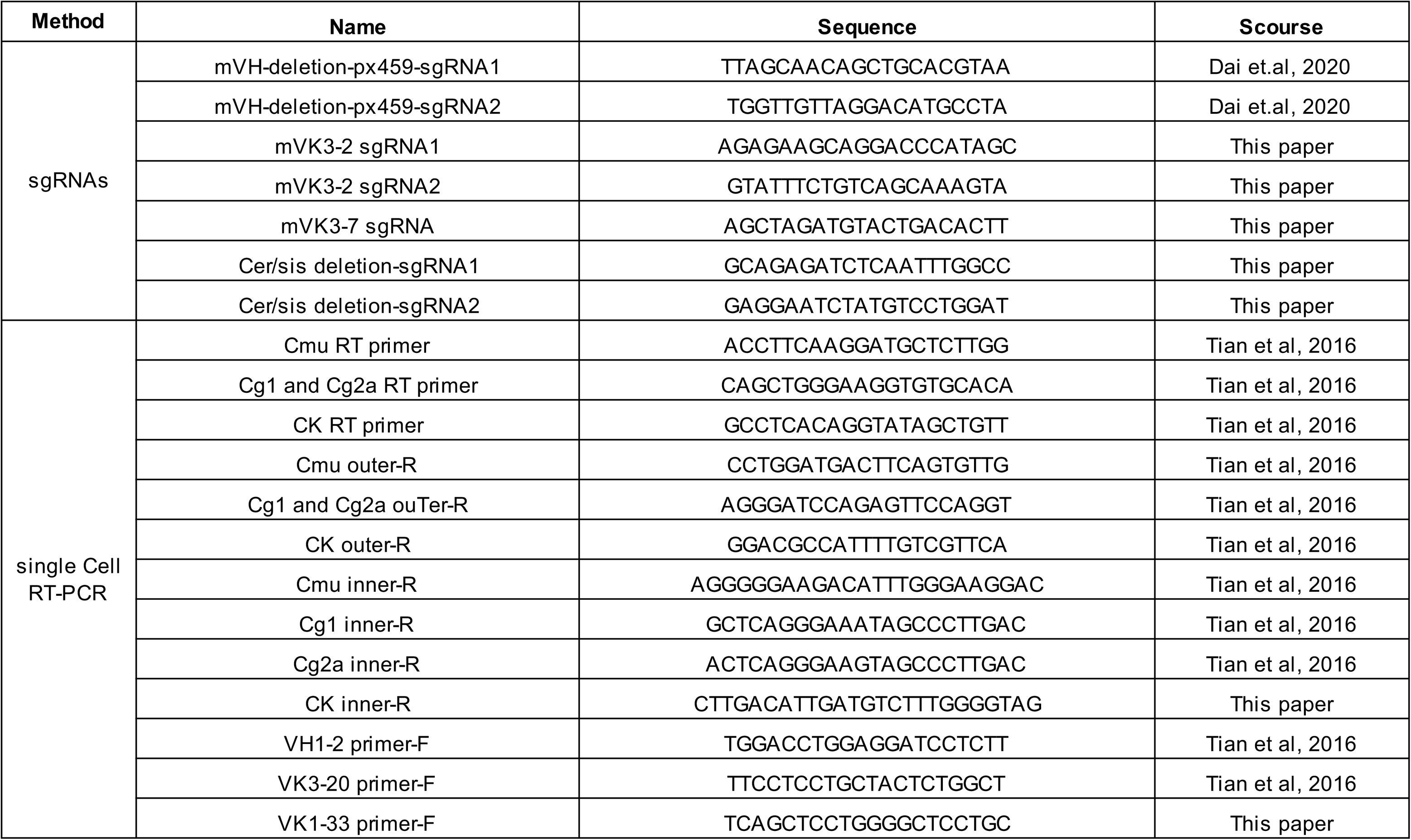
sgRNA sequences for genome editing and primer sequences for single cell RT-PCR.

